# Quality assessment and refinement of chromatin accessibility data using a sequence-based predictive model

**DOI:** 10.1101/2022.02.24.481844

**Authors:** Seong Kyu Han, Yoshiharu Muto, Parker C. Wilson, Aravinda Chakravarti, Benjamin D. Humphreys, Matthew G. Sampson, Dongwon Lee

**Affiliations:** Department of Pediatrics, Division of Nephrology, Boston Children’s Hospital, Boston & Harvard Medical School, Boston, MA, USA; Division of Nephrology, Department of Medicine, Washington University in St. Louis, St. Louis, MO, USA; Department of Pathology and Immunology, Washington University in St. Louis, St. Louis, MO, USA; Center for Human Genetics and Genomics, New York University Grossman School of Medicine, New York, NY, USA; Department of Developmental Biology, Washington University in St. Louis, St. Louis, MO, USA; Broad Institute of MIT and Harvard, Cambridge, MA, USA

## Abstract

Chromatin accessibility assays are central to the genome-wide identification of gene regulatory elements associated with transcriptional regulation. However, the data have highly variable quality arising from several biological and technical factors. To surmount this problem, we use the predictability of open-chromatin peaks from DNA sequence-based machine-learning models to evaluate and refine chromatin accessibility data. Our framework, <u>g</u>apped *<u>k</u>*-<u>m</u>er SVM <u>q</u>uality <u>c</u>heck (gkmQC), provides the quality metrics for a sample based on the prediction accuracy of the trained models. We tested 886 samples with DNase-seq from the ENCODE/Roadmap projects to demonstrate that gkmQC can effectively identify high-quality samples underperforming owing to marginal read depths. Peaks identified in high-quality samples by gkmQC are more accurately aligned at functional regulatory elements, show greater enrichment of regulatory elements harboring functional variants from genome-wide association studies (GWAS), and explain greater heritability of phenotypes from their relevant tissues. Moreover, gkmQC can optimize the peak-calling threshold to identify additional peaks, especially for single-cell chromatin accessibility data as well as bulk data. Here we provide a standalone open-source toolkit (https://github.com/Dongwon-Lee/gkmQC) for such analyses and share improved regulatory maps using gkmQC. These resources will contribute to the functional interpretation of disease-associated regulatory genetic variation.

## Introduction

Open chromatin at specific genomic sites is the hallmark of *cis*-regulatory element (CRE) activity that modulates transcription of a target gene^1^. Thus, identifying open-chromatin regions of the genome is a fundamental step towards defining the gene regulatory program encoded in the genome. Significant advances in experimental techniques to detect these regions have been made over the last decade. Sequencing-based assays, such as ATAC-seq^2^ and DNase-seq^3,4^, detecting transposase-accessible and DNase-hypersensitive regions as open-chromatin regions, are now widely used to enable genome-wide mapping of regulatory elements^5^. These assays have shown that epigenetic landscapes are dynamic and actively regulated across different biological states, cell types^6^, developmental stages^7^, aging^8^, and species^9^. Moreover, multi-omic analyses integrating these data with genome-wide association studies (GWAS) are now significantly accelerating mechanistic understanding of how non-coding variation drives transcriptional regulation^10–12^, the largest contributor to complex traits.

Defining the regulatory landscape requires rigorous assessment of the quality of chromatin accessibility data and this remains a challenge due to several reasons, such as the lack of gold-standard data sets for benchmarking and difficulties in functional validation. Several methods proposed to rectify this include quantifying quality using statistics like the fraction of reads in peaks (FRiP)^13^, signal portion of tags 2 (SPOT2)^14^, and promoter (TSS) enrichment scores,^15^ measuring the degree to which reads are enriched in functional elements (i.e., identified open-chromatin peaks and promoters). Irreproducible discovery rate (IDR) is yet another quality assessment statistic, measuring the reproducibility of peaks between replicates^16^. However, these metrics, such as FRiP and SPOT2, may not be optimal for samples with low sequencing depth where a smaller number of peaks are detected^17,18^. This is problematic, especially for single-cell analysis or rare cell types. IDR is also limited when robust replicates are unavailable. Consequently, chromatin accessibility data with suboptimal quality currently can mislead downstream analyses.

As an improvement, we developed a complementary and biologically motivated quality metric for chromatin accessibility data based solely on their underlying DNA sequences. This metric is based on the concept that CREs, such as promoters, enhancers, and insulators, typically have multiple transcription factor binding sites (TFBSs)^19^. Thus, open-chromatin peaks in high-quality samples (those containing high-quality open-chromatin peaks) are likely to harbor such TFBSs, which can be accurately captured by sequence-based predictive models^20–24^. This leads to our main hypothesis that the accuracy of a sequence-based model directly correlates with the quality of the peaks derived from chromatin accessibility data. Our new method, called <u>g</u>apped *<u>k</u>*-<u>m</u>er SVM <u>q</u>uality <u>c</u>heck (gkmQC), is based on a sequence-based machine learning technique, gkm-SVM^21,25^, which can predict CREs using their primary DNA sequence only. We demonstrate that “high-quality” samples defined by gkmQC (1) have more CREs better aligned at functional regulatory elements, (2) harbor more putatively functional variants from GWAS, (3) and explain greater heritability of traits from their relevant tissues than low-quality samples. We also show that gkmQC can identify additional peaks by optimizing a peak calling process, especially for rare cell types in single-cell chromatin accessibility data.

## Results

### Open-chromatin peak signals correlate with sequence-based prediction performance

A typical chromatin-accessibility data exhibits a wide range of peak signals (i.e., abundance of mapped reads or peak heights). We hypothesized that a peak with a stronger signal, or a higher peak, is more likely to be a true CRE, harboring clearer predictive sequence features, and leading to the higher classification accuracy of sequencebased predictive models. To test this, we analyzed 886 samples of ENCODE DNase-seq^26^. For each sample, we first divided the entire set of peaks, stratified by peak signal strength, into subsets comprised of an equal number of peaks (5,000). We call these “peak subsets.” We then trained gkm-SVM^25^ for each peak subset against an equal number of random genomic regions and performed 5-fold cross-validation to calculate the area under the ROC curve (AUC; i.e., peak predictability). Consistent with our hypotheses, we found a strong correlation between the AUC and peak signals for almost all datasets (**Fig. S1**). Based on this, we defined an overall quality score for a sample as the average AUC score over its degradation rate across peak subsets, dubbed “gkmQC (sample) score” **(Fig. 1; Methods and Materials)**.

**Figure 1.**
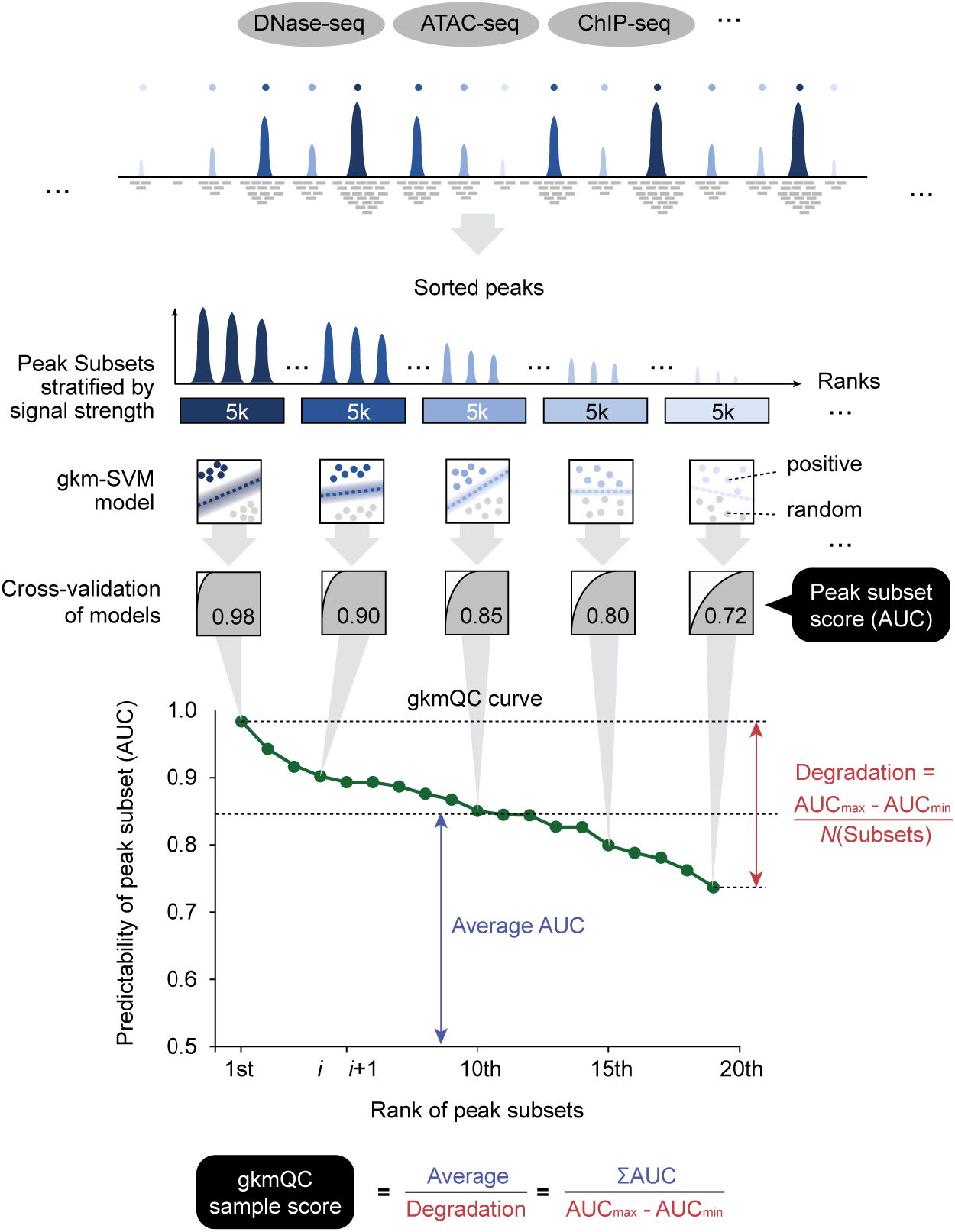
A schematic of peak quality assessment in gkmQC. gkmQC sorts peaks by their signal strengths, groups them into subsets of an equal number of peaks, and calculates the area under the ROC curves (AUCs) using sequencebased predictive models (gkm-SVM) with cross-validation. The AUC represents the overall predictability of peaks in a subset. For a typical data set, AUCs strongly correlate with their peak signal strengths. The curve of AUC scores across subsets reflects its overall sample quality, and the extent of the degradation and the average of AUC across bins is represented by gkmQC sample score.

### Peak predictability complements conventional methods of quality assessment

We next evaluated if the gkmQC score could be used as an alternative sample quality metric. We reasoned that high-quality samples would have greater gkmQC scores as more high-quality peaks in the samples lead to high AUCs and slower degradation of AUC across peak subsets. Using the ENCODE DNase-seq dataset again, we systematically compared their gkmQC scores with SPOT2 scores, a standard DNase-seq quality metric used in the ENCODE project (**Methods and materials;** discussed below) and achieved a significant correlation (*ρ* = 0.66; *P* < 4.9×10^−120^; quality-rank correlation; **Fig. 2a**). We also found several outliers in the correlation (top-left of the scatter plot). To investigate these outliers further, we compared both metrics with two additional quality parameters (**Fig. 2b**). We first analyzed peak coverages, or the number of peaks (|*P*|), and found that both quality metrics (SPOT2 and gkmQC) showed significant correlation (*ρ*_gkmQC_ = 0.49 vs. *ρ*_SPOT2_ = 0.63; quality-rank correlation; **Fig. 2c**). We next evaluated how well peaks are aligned with known regulatory elements (i.e., precision of peak location). Specifically, we quantified genomic distance between the peak summit locations and the center of overlapping FANTOM5 enhancers^27^, which are known to be strong and robust distal regulatory elements in the human genome (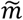; **Methods and materials**). We found that gkmQC score strongly correlated (*ρ* = 0.53) with 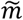 while SPOT2 showed only marginal correlation (*ρ* = 0.23), suggesting that gkmQC can identify high-quality samples defined by those peaks with a precise location for their functional regulatory elements (**Fig. 2c**). We note that these two quality parameters are independent as the correlation of 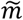 and |*P*| is −0.21 across the ENCODE DNase-seq samples (**Fig. S2**). We also assessed whether other technical conditions, such as sample types, treatments, and sequencing steps, affected these quality metrics and confirmed that no other confounding factors significantly affected gkmQC score and other metrics (**Fig. S3**). In sum, our results suggest that gkmQC can reclassify some low-quality samples as determined by SPOT2 into high-quality samples.

**Figure 2.**
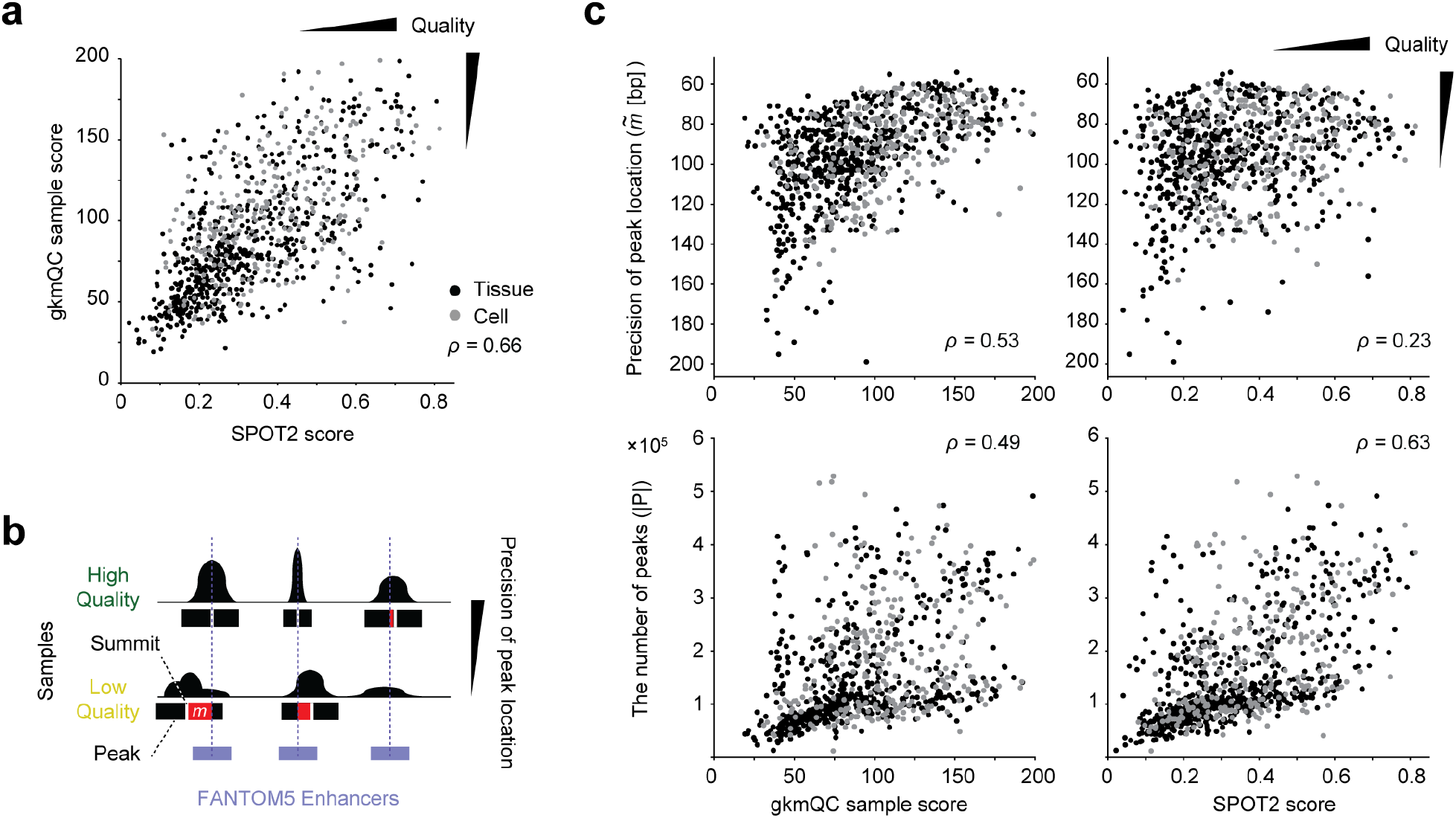
Quality parameters of peaks assessed by gkmQC and SPOT2. **(a)** A scatter plot of gkmQC score and SPOT2. **(b)** A concept diagram depicts mapping precisions of peaks in high- and low-quality samples. The red bar shows *m* that represents the positional mismatch of a peak and the overlapping FANTOM5 enhancer. **(c)** The two quality metrics (X-axis; gkmQC sample score and SPOT2) are compared to the two parameters (Y-axis; locating precision and the number of peaks) in the scatter plots. The Y-axis for the *m* parameter is reversed so that samples in right-top corners are high-quality samples. Spearman’s rank correlation coefficients (*ρ*) are calculated for these comparisons. Black and gray dots are tissues and cells, respectively.

### High-quality samples reproduce precise locations of peaks relevant to GWAS variants

Accurate identification of regulatory elements is crucial to interpreting changes in the regulatory functions of relevant GWAS variants^28^. We hypothesized that open-chromatin peaks that contain functional regulatory variants are more precisely identified in high-than low-quality samples (**Fig. 3a**). We assumed that the consensus peak summit (i.e., centroid) of open chromatin peaks across multiple replicates represents the core functional regulatory element as previously shown in Meuleman et al^29^. We then tested if peaks in high-quality samples are in greater proximity to these centroids. We first focused on a few loci with putative causal GWAS variants for a major kidney functional trait, estimated glomerular filtration rate (eGFR)^30^, using developing kidney DNase-seq samples^26^. For example, rs11261022 is a likely causal regulatory variant affecting eGFR via change of *KLHDC7A* expression^30^. This variant is in an open-chromatin peak in kidney samples (**Fig. 3b**). However, most lower quality samples either do not have an overlapping peak or have a peak less optimally aligned with the variant. Similarly, rs77924615 is another putative functional regulatory variant identified by fine-mapping and colocalization analysis. Again, while high-quality samples have an open-chromatin peak well aligned with this variant, low-quality samples failed to detect open-chromatin peaks in this locus (**Fig. 3c**). Moreover, peaks in high-quality samples exhibit stronger signals than low-quality ones, demonstrating their greater utility in identifying regulatory elements containing GWAS variants (**Fig. 3b and c**).

**Figure 3.**
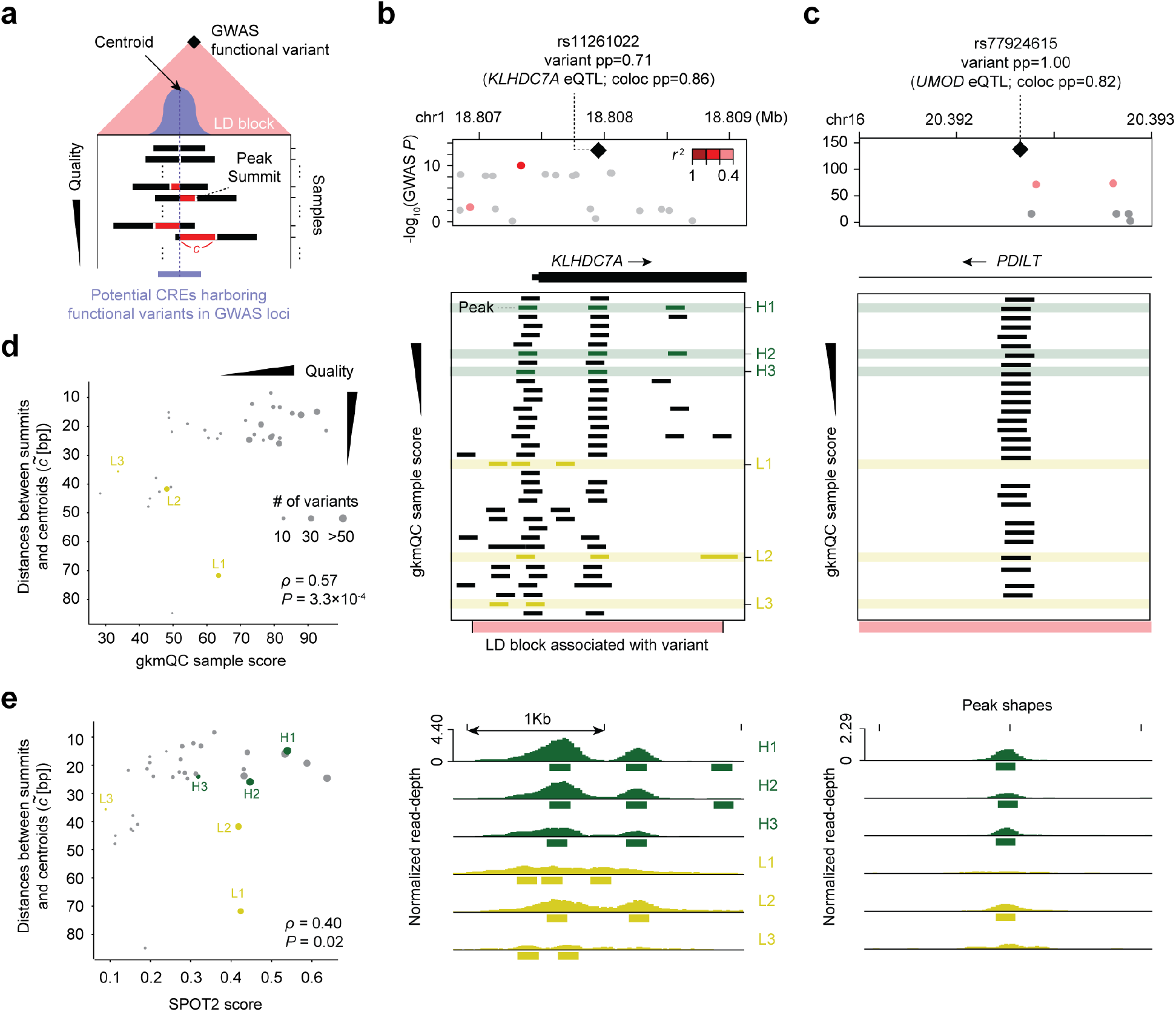
Peaks in high-quality samples are better aligned with the centroid of peaks across biological replicates near GWAS functional variants. **(a)** The centroid of peaks across replicates represents the hypothetical core of a functional CRE (light purple bar). Peak summits in high-quality samples are better aligned with their centroids across replicates than the low-quality ones. Each bar shows a peak from the corresponding sample. *c* (red-colored bar) is the genomic distance between a peak summit and its centroid. **(b-c)** Representative examples of peaks associated with functional GWAS variants in the **(b)** *KLHDC7A* and **(c)** *PDILT* locus. Rows are samples sorted by gkmQC scores, so top samples are high-quality. The bottom panels are the read pileup visualizations for the peak shapes shown in top panels. **(d-e)** gkmQC and SPOT2 scores are compared with 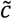 for all kidney samples. H1-3 and L1-3 samples are representative examples of high- and low-quality samples classified by gkmQC, respectively. Each dot represents a developing kidney sample. The dot size corresponds to the number of GWAS loci harboring functional variants within <1kb from the peaks in relevant samples.

To generalize this finding, we calculated genomic distances between the peak summits from each sample and the centroids of the consensus peaks near each putative causal GWAS variant (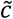; **Methods and materials**). We found that high-quality samples determined by gkmQC have smaller average distances than low-quality samples. Overall, gkmQC score strongly correlated with 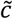 (*ρ* = 0.57; *P* = 3.3×10^−4^; quality-rank correlation; **Fig. 3d**), while conventional metrics, such as SPOT2, were ambiguous (*ρ* = 0.40, *P* = 0.02; **Fig. 3e**). Thus, peaks in high-quality samples determined by gkmQC more precisely identify functional regulatory elements with functional GWAS variants.

### High-quality samples systematically improve the discovery of the genetic architecture of complex traits

The above examples suggested that high-quality samples could improve the systematic identification of genomewide polygenic signals. To test this, we adapted the stratified LD-score regression (S-LDSC) and performed partitioning heritability analyses of many human traits. Specifically, we calculated a normalized S-LDSC coefficient, which corresponds to a statistical significance of the per-SNP heritability (**Methods and materials**)^31,32^. Using replicate kidney samples during development, we found that gkmQC score significantly correlated with S-LDSC coefficients for the eGFR (*ρ* = 0.76, *P* = 1.3×10^−7^; Pearson correlation coefficient; **Fig. 4a**)^30^. This suggests that the gkmQC scores for sample quality significantly explain the variance in heritability across biological replicates. Next, we extended our analysis to multiple tissues and traits (**Methods and materials**). Using 214 samples from 8 tissues/cells from the ENCODE project and 30 different traits from the UK-Biobank^33^, we discovered that high-quality samples stratified by gkmQC consistently gave higher heritability signals, especially for relevant trait-tissue combinations (**Fig. 4b; Fig. S4 and 5a and b**). Most samples with gkmQC scores >70 achieved S-LDSC *z* >1.0 for their relevant traits (**Fig. S4**), suggesting that this is a reasonable threshold for identifying good quality samples. We also achieved comparable results with SPOT2 scores (**Fig. S5c**), demonstrating that variation in S-LDSC coefficients across samples from the same tissues are explained by sample quality factored by peak coverage (**Fig. 2c**). We note that S-LDSC relies on an LD reference panel to estimate heritability based on haplotype blocks (>1kb)^32,34^. Therefore, it may not explain the variation of sample quality factored by precise peak locations (<100bp; **Fig. 3d and e**), which only gkmQC can explain. Taken together, our results suggest that functionally important tissuespecific regulatory elements are better detected in high-quality samples.

**Figure 4.**
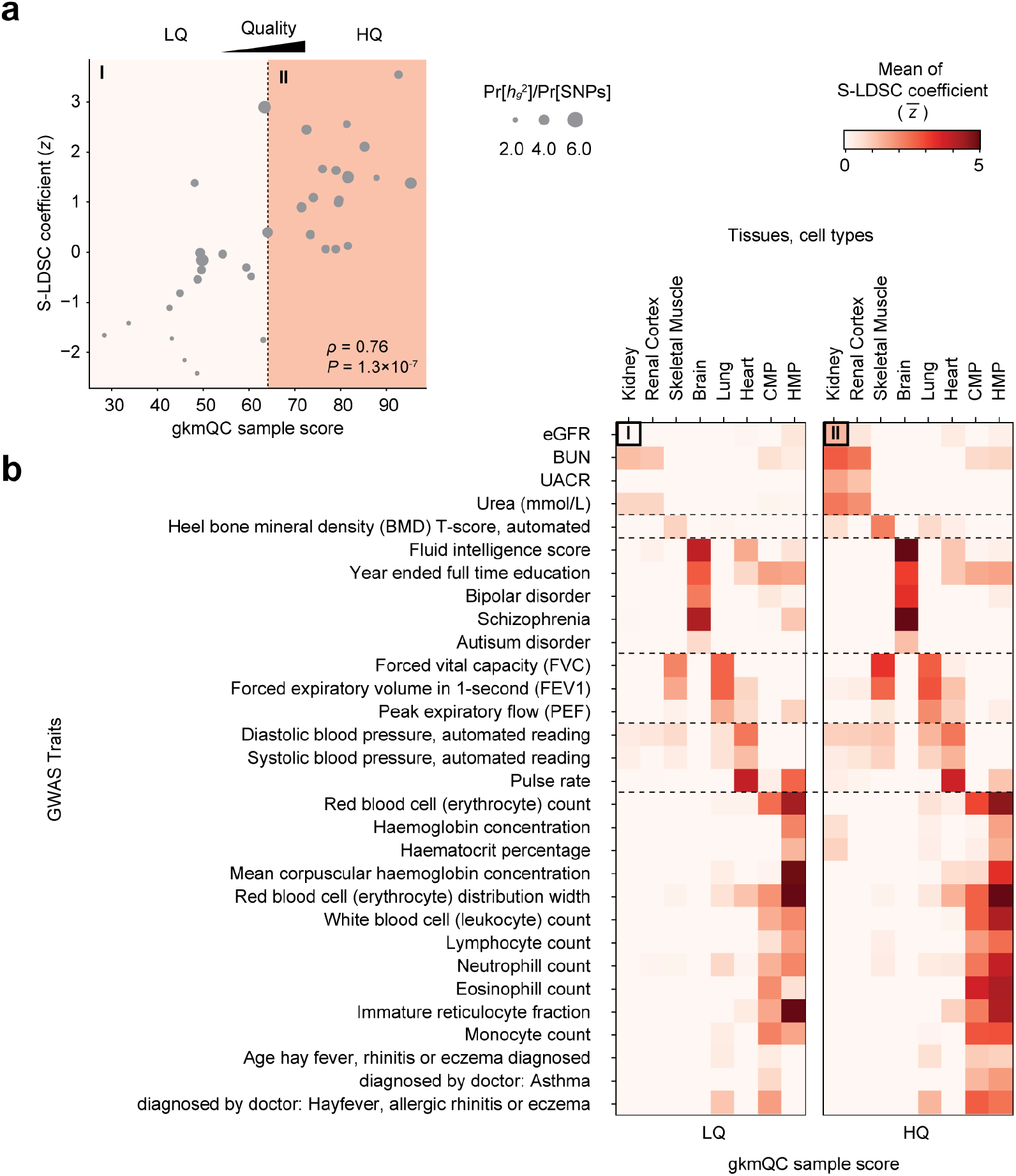
Peaks in high-quality samples exhibit greater heritability for relevant phenotypes. **(a)** A scatter plot comparing gkmQC score and normalized S-LDSC coefficient (z-score) for eGFR is shown for 35 developing-kidney samples. The S-LDSC coefficient directly correlates with the enrichment score of heritability for eGFR. **(b)** Two heatmaps comparing the average S-LDSC coefficients between high- and low-quality samples. We used the top 50% gkmQC score*s* as a threshold for sample quality classification. The top-left cells of the two heatmaps summarize the scatterplot of (a). CMP and HMP are the abbreviation of common myeloid progenitor and hematopoietic multipotent myeloid progenitor cells, respectively.

### gkmQC optimizes the sensitivity on peak-calling

A benefit of gkmQC is that its peak predictability can be used for recovering peaks that do not pass the statistical threshold based on read enrichment but have strong sequence signatures nonetheless. While analyzing the full set of ENCODE data, we noticed that 58 samples with lower read depth exhibited high AUC for all subsets in the sample (i.e., Minimum AUC [MinAUC] >0.75; **Fig. S6a**). This result suggested that additional CREs with strong sequence signatures were undetected. We reasoned that this might be due to an overly stringent peak calling threshold given low read depths. Our approach can, however, naturally alleviate this issue by identifying a more accurate threshold based on peak predictability. To test this, we recalled peaks using a conventional peak caller but with a relaxed threshold and then applied gkmQC to find a new peak calling threshold that maximizes the number of peaks with MinAUC >0.7 (**Fig. 5a-b; Methods and materials**).

**Figure 5.**
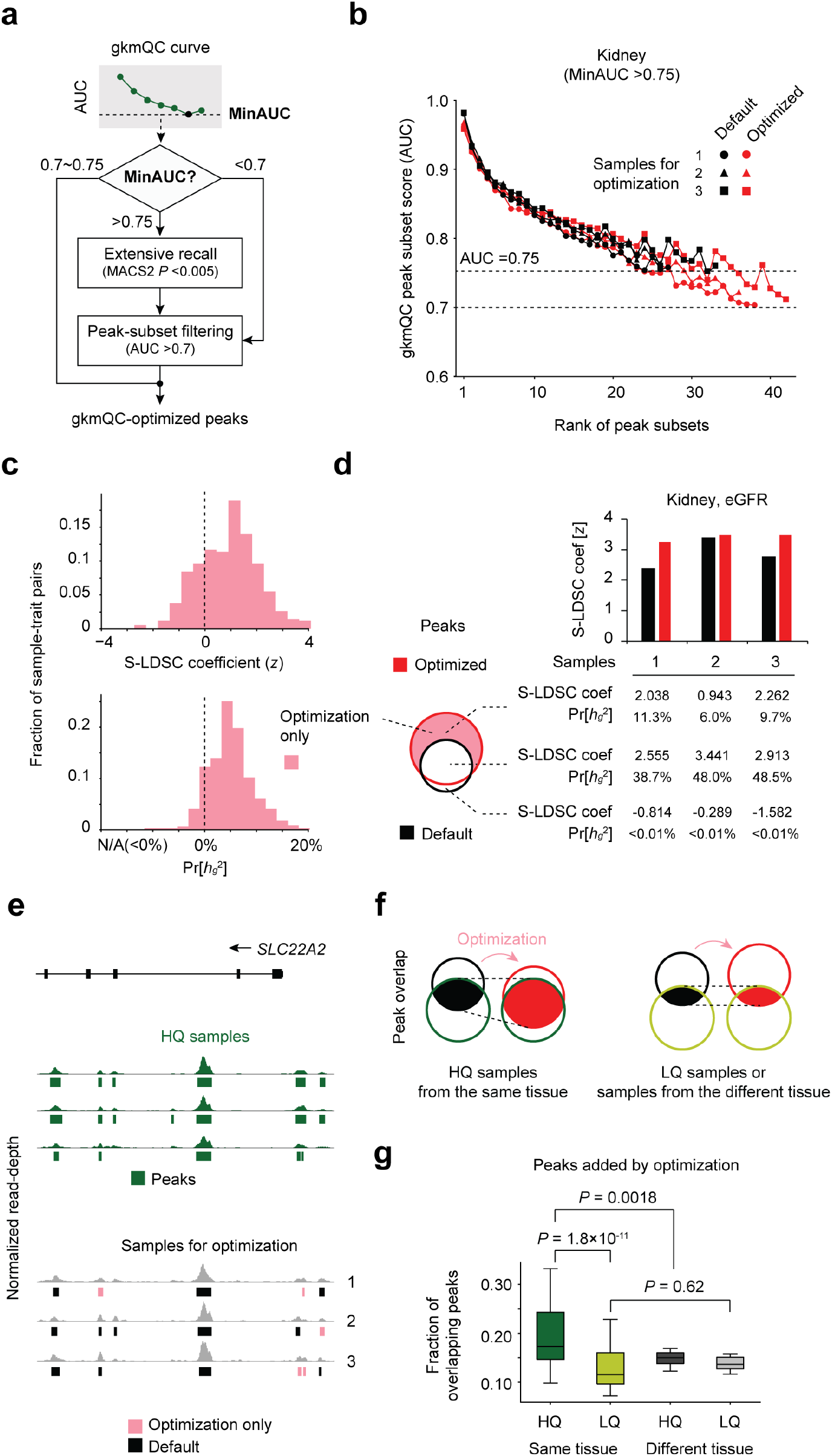
Optimization of peak calling in bulk chromatin accessibility data. **(a)** Workflow diagram of peak-calling optimization process using gkmQC. **(b)** gkmQC curves for three representative developing-kidney samples before (Black) and after optimization (Red). **(c)** Distribution of S-LDSC coefficient scores and the proportion of the heritability of newly identified peaks after optimization (i.e., optimization-only; pink in the Venn diagram). Heritability was calculated for 58 samples in 8 tissues and 31 relevant phenotypes requiring optimization. **(d)** Heritability analysis for three representative developing-kidney samples. The graphs of black and red bars show the S-LDSC coefficients calculated with whole peaks before and after optimization. The table presents heritability for three subsets of peaks; optimization-only (top), commonly found regardless of the optimization (middle), and with default parameters only (bottom). **(e)** A specific example of open chromatin peaks in the *SLC22A2* locus. Green bars represent peaks from high-quality samples. Dark grey and pink bars are the commonly found and optimization-only peaks in the samples targeted for the optimization step. **(f)** A conceptual diagram to show that gkmQC optimization increases the reproducibility of peaks in high-quality samples from the same tissue. **(g)** Comparisons of overlaps between peaks added by optimization and those in other samples. Box plots show the fraction of peaks recovered by gkmQC optimization overlapping those in high-quality (HQ) replicates (green), low-quality (LQ) replicates (light green), HQ samples from different tissues (black), and LQ samples from different tissues (gray). Paired *t*-test is used to test the significance of the differences between these groups.

Among the DNase-seq data of bulk tissues and cells, 58 of 200 samples with low read-depths had MinAUC >0.75. gkmQC optimization of these samples recovered an additional ^~^27.2% of total peaks per sample on average (**Fig. S6b**). To check the functional relevance of recovered peaks, we repeated the S-LDSC analyses again for traits relevant to the samples. To account for potentially inflated heritability caused by SNPs in strong LD, we analyzed the newly identified peaks in S-LDSC while controlling for default peaks (**Methods and materials**). We achieved positive S-LDSC coefficients in 76.4% of all datasets and increased proportion of the heritability for 89.2% of the data sets, suggesting that the newly identified peaks by gkmQC were trait-relevant (**Fig. 5c**). We also observed that the optimized versus default peaks achieved consistently higher heritability (**Fig. S6c**). In **Fig 5e**, we provide a specific example of newly identified peaks at the kidney-specific gene, *SLC22A2* locus. Here, we focused on three kidney samples with MinAUC >0.75, in which additionally discovered peaks with AUC >0.7 produced positive S-LDSC coefficients (**Fig. 5b-c**). Significantly, recovered peaks at this locus (pink bars) overlapped those identified in high-quality samples of the kidney (**Fig. 5e and Fig. S6d**; *KLHDC7A*).

Inspired by the cases of *KLHDC7A* and *SLC22A2*, we next hypothesized that gkmQC optimization could improve the reproducibility of peaks given the quality variation across replicates (**Fig. 5f**). To validate this systematically, we investigated if the newly found peaks are replicated in other high-quality samples with the same tissue entity. Specifically, we calculated the fraction of the newly found peaks by optimization that overlaps with peaks in high-quality samples from the same tissue (**Methods and materials**). We used low-quality samples from the same tissue and high-quality samples from different tissues as negative controls for comparison. We found that the peak overlaps with high-quality replicates significantly higher than low-quality replicates (**Fig. 5g;** *P* = 1.8×10^−11^; Paired *t*-test) and high-quality samples from different tissues (*P* = 0.0018). Thus, our gkmQC optimization can considerably recover tissue-relevant peaks that are present in high-quality samples.

### Extensive peak calling by gkmQC improves single-nucleus ATAC-seq inference

Because gkmQC successfully optimized bulk data with marginal read-depth (**Fig. 5**), we surmised that gkmQC could also be helpful in improving peak identification in single cell chromatin accessibility data, especially for rare cell types, which by nature have a lower numbers of reads^35^. To test this, we applied the peak optimization process to kidney single-nucleus ATAC-seq data (snATAC-seq)^36^. We first analyzed a rare but important kidney cell type of the glomerular filtration barrier, the podocyte (<1% of total kidney cells) (**Fig. 6a and Fig. S7a-d; Methods and materials**)^36^. Our gkmQC peak calling optimization recovered ^~^35,000 additional peaks in podocytes compared to peaks called with a default threshold. Second, the variants in these recovered peaks significantly contributed to the heritability of Urine Albumin-to-Creatinine Ratio (UACR; *z* = 3.357, S-LDSC coefficient; **Fig. 6b**), a kidney trait affected by podocytes. In contrast, no significant contribution to the heritability of unrelated traits such as schizophrenia^37^ was observed (SCZ; *z* = 0.010). **Fig. 6c** shows a specific example of peaks recovered at the podocyte-relevant *LMX1B* locus, a well-known podocyte-specific transcription factor for kidney development and the maintenance of differentiated podocytes^38,39^. We note that a recovered peak upstream of *LMX1B* was also detected in bulk developing kidney tissues (**Fig. 6c**). Since this peak was not detected in any other kidney cell types (**Fig. S7e**), it demonstrates that optimization using gkmQC can uncover new peaks even in rare cell types, which are more likely to have peaks missed because of their lower numbers.

**Figure 6.**
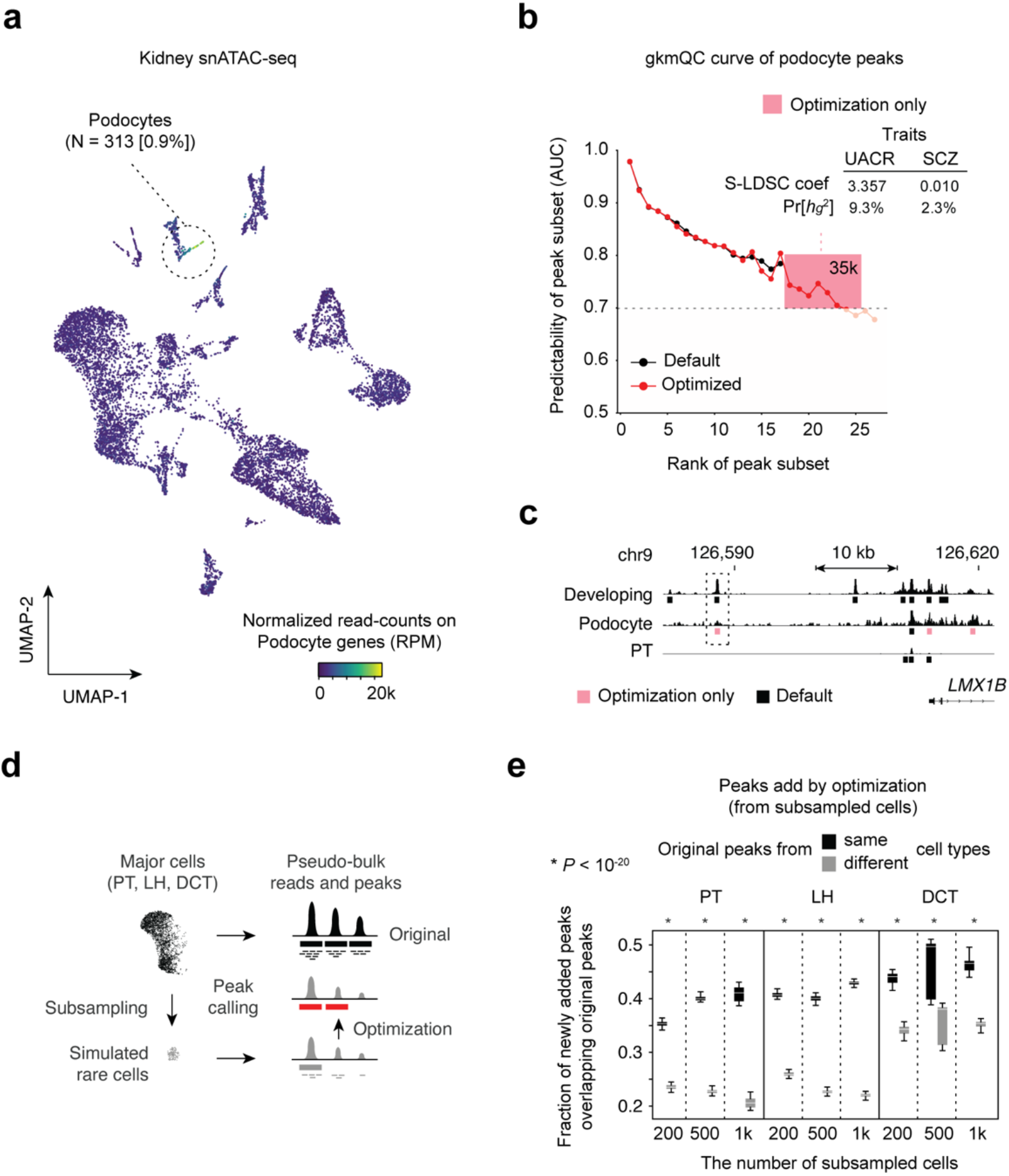
Improvement of peak-calling in snATAC-seq data for rare cell types. **(a)** A UMAP plot of kidney snATAC-seq analysis. The heatmap colors represent open-chromatin activities of podocyte-specific genes. **(b)** gkmQC curves for pseudo-bulk reads of podocyte cells are shown before and after optimization. Peak subsets within the pink box are the ones newly discovered by optimization. The inset table shows a heritability analysis result for the podocyte-relevant (Urine albumin-to-creatinine ratio; UACR) and one non-relevant (Schizophrenia; SCZ) trait for these newly identified peaks only **(c)** A specific example of podocyte open chromatin peaks in the *LMX1B* locus. The pink bar is a newly called peak by optimization. PT is the proximal tubule, the most abundant cell type in the kidney, and is used as a control to show cell-type specificity of the new peaks. **(d)** A schematic describing how gkmQC optimization is assessed using random subsampling of major cell types to simulate rare cell types. **(e)** Box plots represent fractions of optimized peaks in subsampled cells overlapping the original peaks. LH and DCT are the abbreviation of loop of Henle and distal convolute tubule cells, respectively (**Fig. S8**).

To systematically test if the optimization process is generalizable to other cell-types and tissues, we analyzed the heritability contributed by variants in recovered peaks from the other 11 kidney cell-types and peripheral blood mononuclear cells (PBMC) (**Fig. S8**). We discovered that, in general, cell-types with lower cell counts had higher MinAUCs (**Fig. S7d and Fig. S8d**), showing that the optimization process is more effective in rarer cell-types. Expectedly, we confirmed that the newly identified peaks in rare cell types with counts < 1,000 explained a significant proportion of heritability (S-LDSC coefficient >2.0) for relevant traits (**Fig. S7f and S7e**).

Lastly, we subsampled 200, 500, and 1,000 cells from five abundant cell types from kidney and PBMC (i.e., Proximal Tubule (PT), Loop of Henle (LH), and Distal Convoluted Tubule (DCT) for kidney, and CD4^+^ Memory T cells (CD4M) and CD14^+^ Monocytes (CD14M) for PBMC) to simulate rare cell populations and to test if gkmQC peak calling optimization for sub-sampled cells can recapitulate peaks identified in the original populations of the cells (**Methods and materials**). We found that 30^~^40% of peaks added by gkmQC optimization overlapped the original peaks from the corresponding cell types (**Fig. 6d-e and Fig. S8f**). The fraction of the overlapped peaks was significantly higher than that of original peaks from different cell types (*P* < 10^−20^; Paired *t*-test). Taken together, these results show that the peak-calling optimization using gkmQC helps us to identify additional cell-type-specific regulatory elements, particularly for rare cell types.

## Discussion

Comprehensive quality control (QC) analysis of chromatin accessibility data is a critical need for proper analysis of genome regulatory functions because of lack of ground-truth datasets for benchmarking. By utilizing machinelearning techniques which learn sequence features underlying open-chromatin peaks, we have established a computational framework for quality assessment and refinement of chromatin accessibility data. High-quality samples determined by gkmQC yield more accurate data for more robust downstream genomic analyses, such as GWAS fine-mapping of complex traits and partitioning its heritability. Our method for optimizing peak calling thresholds also improves single-cell chromatin accessibility datasets by identifying more peaks in rare cell types.

Precise mapping of functional regulatory elements is now possible by applying several genomic technologies^40–42^. A recent study demonstrated that the centroid of overlapping consensus DNase I Hypersensitive site (DHS) summits can be used to robustly and accurately identify core regulatory regions in which variants that disrupt TF bindings strongly perturb their regulatory activities^29^. We show that peaks in high-quality samples identified by gkmQC can also accurately locate these core regulatory regions (**Fig. 3**). Highly predictive sequence features presented in peaks in high-quality samples enable us to precisely locate these elements, while quality control using these sequence features can be used to test the precision of peak location without many replicates.

Peak subsets with medium-level AUCs (0.85-0.95) are enriched for distal enhancers and tissue-specific regulatory elements (**Fig. S9 and S10)**. In contrast, peak subsets with high AUCs (>0.95) mostly contain promoters and ubiquitously open regions with homogeneous sequence features. These results imply that tissue-specific CREs and distal enhancers may have consistently lower peak predictability than housekeeping ones and promoters regardless of their sample qualities. It is also concordant with the known biology that regulatory activity of these tissue-specific and distal enhancers could be modulated by changes in TF expression or chromatin looping without strong sequence-specific binding of TFs^43^. Thus, our strategy of evaluating multiple peak subsets stratified by signal strength independently, rather than building a single model on the whole peak set, makes it possible to assess sample qualities more accurately by allowing multiple models to capture different classes of CREs active in a sample.

Comprehensive identification of peaks for rare cell types is a big challenge in single-cell ATAC-seq analysis^35^. We showed that our gkmQC optimization could find ^~^27% more peaks in rare cell types (**Fig. 6 and S7-8**). It also enables us to estimate the minimum number of cells required for adequate peak identification, which is currently unknown. We have found that, in general, cell types with cell counts >1,000 can yield >100,000 peaks and do not need gkmQC optimization in general (**Fig. S7b and S8b**). Considering that >100,000 peaks are typically identified in bulk DNase-seq and ATAC-seq from cell lines and primary cells, we speculate that at least 1,000 cells are needed for a comprehensive peak discovery. We also note that optimizing cell clustering and peak-calling algorithms are required for achieving better peak discovery.

Chromatin accessibility data have empowered us to functionally interpret disease-associated genetic variation. Over the last decade, a significant amount of chromatin accessibility data has been accumulated to create an atlas of functional regulatory elements across diverse tissues or cells. We anticipate that our new quality assessment framework will further accelerate this process by prioritizing high-quality samples, finding more CREs from rare cell types in snATAC-seq, and implicating sequence variants that disrupt these functions.

## Methods and Materials

### Sequence-based predictive model for quality evaluation

We constructed gkm-SVM models following our previously established framework^25,44^. Briefly, we first defined open-chromatin regions derived from a DNase-seq dataset as the positive training set using a precalculated openchromatin peak set available in ENCODE^26^. A negative set for training was then generated by random sampling of an equal number of genomic regions that match the length, GC content, and repeat fraction of the positive set. We excluded regions with >1% N-bases and >70% repeats from the training datasets. To prevent potential bias caused by variable peak lengths, we fixed the size of peaks by extending 300bp from the summit.

We split the genomic peaks into multiple subsets, each comprising 5,000 peaks sorted by decreasing signal intensity scores from a peak caller. If a group of peaks with the same score were separated into two subsets, we randomized the order of these peaks to make sure that all neighboring peaks (i.e., peaks sorted by genomic position) were not grouped into the same subset. Then, we trained a gkm-SVM model for each peak subset using default parameters (word length *l* = 10, informative columns *k* = 6, truncated filter *d* = 3, and weighted gkm kernel (wgkm) *t* = 4). Model performance was measured by the area under the ROC curves with five-fold cross-validation (i.e., an AUC-score of the peak subset). We present AUC scores and ranks of peak subsets as a gkmQC curve on the Y- and X-axis, respectively.

To quantify overall sample quality, we derived a gkmQC score from a gkmQC curve using the following equation,

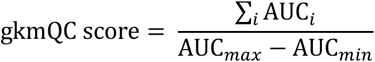

where *i* is the rank-numbers of peak subsets and AUCmax and AUCmin are the maximum and minimum value of AUC scores in the curve, respectively. We limit analysis to the top 100,000 peaks (= 20 subsets of 5,000 peaks per subset) to generate a gkmQC curve. We did so to reduce the computation cost based on the observation that most ENCODE samples have ^~^100,000 peaks.

To analyze sequence features of the models, we scored all possible 10-mers (*l* = 10) using the trained SVM model and then performed a principal component analysis (PCA) using their score vectors. We evaluated the first two principal components to compare models (**Fig. S9d and e**). To analyze tissue specificity of peak subsets, we calculated pairwise peak overlaps between the same rank subsets from the same tissues. To ensure these peak overlaps were comparable across different rank peak subsets, we further normalized them by calculating an overlap fold-change for each rank. As a denominator, we used the average overlap between the same rank peak subsets from random tissue pairs.

### Benchmarking datasets and evaluation methods

#### ENCODE DNase-seq datasets

We revisited the ENCODE DNase-seq datasets accessed on 03/10/20. We excluded samples with <10,000 peaks, archived, or revoked status in the database. We ultimately obtained 886 DNase-seq datasets across diverse samples, including *in vitro* differentiated, primary cells, and tissues. Metadata of the full datasets are in **Table S1**.

#### Quality parameters for peaks

To obtain the precision of peak location 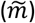, we calculated the average genomic distance between peak summits and the center of overlapping FANTOM5 enhancers (https://fantom.gsc.riken.jp/5/datafiles/reprocessed/hg38_latest/extra/F5.hg38.enhancers.bed.gz), identified by Cap analysis gene expression (CAGE)^45^. FANTOM5 enhancers were called based on CAGE peaks with bidirectional balanced RNA signatures, distal to known exons (+/-100bp region from boundaries) and transcription start sites (+/-300bp)^27^. We considered an overlapping enhancer the most proximal enhancer that also overlaps with the 1kb-extended peak from the summit. The peak coverage, |*P*|, is the count of non-overlapping peaks.

We used FANTOM5 enhancers again to analyze the enrichment of open-chromatin peaks for known enhancers. For the enrichment analysis with promoters, we used FANTOM5 promoters (https://fantom.gsc.riken.jp/5/datafiles/reprocessed/hg38_latest/extra/CAGE_peaks/hg38_fair+new_CAGE_peaks_phase1and2.bed.gz). The FANTOM5 promoters were called based on CAGE peaks of which signal is comparable to CAGE peaks near 5’-ends of known transcripts (within 500bp)^46^.

#### Conventional metrics for quality evaluation

To compare our findings with the default metrics provided by ENCODE, we used Signal Portion of Tag (SPOT2), the number of cleavages observed within HOTSPOT2 peaks divided by the total number of cleavages in a sample^14^. We obtained precalculated SPOT2 scores from the metadata of the ENCODE database. We used featureCounts^47^ to calculate the fraction of reads in called-peaks (FRiP), known promoters, and enhancers. Irreproducible discovery rate (IDR) of peak-calling across biological duplicates was calculated by the *IDR* package^16^. To quantify a representative value of IDR values for a sample, we averaged −log10 of IDR *P*-values of all peaks in a sample. To measure the quality affected by sample treatment, especially for autopsied tissues, we curated the duration of postmortem time from cardiac cessation to freezing of the sample from the case report in the ENTEx dataset (https://www.encodeproject.org/entex-matrix/?type=Experiment&status=released&internal_tags=ENTEx)^48^.

### Validation of high-quality samples using GWAS functional variants

#### GWAS datasets

For integrative analyses of open-chromatin peaks with relevant GWAS variants, we focused on GWAS for estimated glomerular filtration rate (eGFR)^30^, a quantitative kidney functional trait. We chose eGFR GWAS due to the significant heritability and the availability of fine-mapping datasets with eQTL co-localization. We specifically used precalculated dataset of putatively functional SNPs from the European-ancestry meta-analysis of the eGFR trait^30^ based on posterior probability >0.5 from approximate Bayes factor analysis^49^.

#### Associating core regulatory elements near functional GWAS variants

We measured the average genomic distance 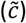 between peak summits in a sample and the centroids of overlapping peaks across biological replicates from the same tissue. We only considered peaks near putatively functional GWAS variants, defined as genomic regions harboring fine-mapped SNPs and their neighboring SNPs in high LD (*r*^2^ >0.8) with a 1,000-base padding. To measure the centroid, we calculated average genomic positions of overlapping peak summits across 35 replicates of developing kidney. Similar to the peaks, we only used the centroids near fine-mapped GWAS variants. We used LocusZoom to plot association *P*-values of GWAS variants with linkage disequilibrium information from a reference population^50^. IGV was used to plot read pile-up signals from open-chromatin data^51^.

### Validation of high-quality samples using partitioned heritability analysis

#### GWAS datasets

We obtained GWAS summary statistics data for various phenotypes from the UK Biobank project^52^ as processed by Neale lab (http://www.nealelab.is/uk-biobank/) and three quantitative kidney traits: eGFR, urine albumin-to-creatinine ratio (UACR) and blood urea nitrogen (BUN)^30,53^. To analyze the relevant GWASs with a significant genetic association, we limited GWASs with heritability *z*-scores >4 (z4 and z7) and medium/high confidence ratings (available in https://nealelab.github.io/UKBB_ldsc/h2_browser.html). We also selected tissues with ≥5 replicates, for which relevant GWASs are available. Consequently, we derived six developing tissues, two primary cells, and 30 relevant GWAS traits (**Table S2**).

#### Stratified LD-score regression

To estimate a proportion of heritability 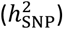 from GWAS summary statistics, we used LD-score regression (LDSC)^31,32^. We employed stratified LDSC to calculate a proportion of heritability contributed to a SNP set (*C*) in open-chromatin peaks from a sample:

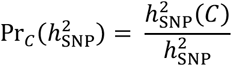

Enrichment of the proportional heritability is presented by 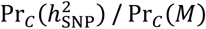, where Pr_*C*_(*M*) is the proportion of SNPs in *C* among the total SNP set. It represents a relative polygenic contribution of SNPs within openchromatin regions to a given trait. To compute a representative parameter reflecting both an effect size and a statistical significance of the enrichment, we used a normalized S-LDSC coefficient driven by,

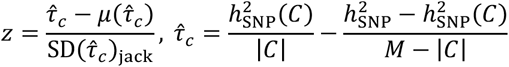

Where 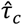 is a normalized coefficient of the proportional heritability to enable *z*-based scoring, and 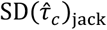 is the standard deviation of 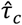 calculated from a jack-knife estimation.

To take into account potential regulatory variants in the flanking regions of open-chromatin peaks, we defined genomic annotations of CREs as a 1kb-padding from the peak summits. To accurately test the contribution of open-chromatin peaks only, we calculated the partitioned heritability along with the baseline annotations of 97 functional regions, such as protein-coding, evolutionary-conserved, promoter, enhancer, or UTR, as recommended by the original S-LDSC study^32^. For reference LD scores, European ancestry population and the corresponding allele frequencies in 1000 Genomes Phase 3 data were used. All data, including the baseline LD annotation set (v2.2), were obtained from https://data.broadinstitute.org/alkesgroup/LDSCORE/. When we compare multiple functional annotations (i.e., optimized versus default open-chromatin peaks), we conducted S-LDSC regression jointly with the multiple annotations, along with the full set of the baseline annotations in one model.

### gkmQC-based optimization of peak-calling threshold

#### The default pipeline for peak-calling

As a default pipeline for peak-calling, we adapted the previously established framework for DNase-seq and ATAC-seq analyses^44,54^. Specifically, we ran MACS2^55^ with no restricted model (--nomodel) using paired-end read pairs with MAPQ >30. We used bp of window-size for calling 100bp (--extsize 100), the −50bp of shifts toward lagging strand (--shift −50), keeping duplicate reads (--keep-dup), and *q*-value cut-off <0.01^56^. For ATAC-seq samples, we additionally trimmed +4bp of the forward strand and −5bp of the reverse strand to account for 9bp duplicated regions by Tn5^2^.

#### gkmQC peak-calling optimization

We first determine if a sample needs a peak-calling optimization based on the minimum AUC of all peak subsets (MinAUC). If the MinAUC is >0.75, we perform the following peak-calling optimization. To recover marginal peaks with suboptimal *q*-values, we called peaks again using a relaxed threshold with nominal *P* <0.005. We then calculated AUCs for *all* peak subsets using our gkmQC framework. Lastly, we recover all peaks that are more significant than the least significant peak in the minimum rank peak subset with AUC >0.7.

#### Analysis of the overlap of peaks added by optimization

To measure peak overlaps between samples, we calculated the fraction of the newly added peaks that overlap with peaks in other samples from the same tissue. We defined high-quality replicates as samples with the top 50% percentile of gkmQC scores among all samples from the corresponding tissue. The rest were used as low-quality replicates. Because the variation of peak counts across samples can be a potential confounding factor for this analysis, we randomly chose 100,000 peaks from a sample. We repeated this process ten times to obtain average overlaps across the ten different realizations of random peak sets.

### Analysis of snATAC-seq data

#### single-cell ATAC-seq datasets

Human kidney snATAC-seq data are from non-tumor kidney cortex samples from five patients undergoing partial or radical nephrectomy^36^. We specifically downloaded sequencing data from GEO under accession number GSE151302. For the dataset of peripheral blood mononuclear cells (PBMC), we used public snATAC-seq data with total ^~^15,000 cells from a healthy donor (Next GEM v1.1). We downloaded position-sorted BAM files derived from Cell Ranger ATAC 1.1.0 from the 10X Genomics support page (https://support.10xgenomics.com/single-cell-atac/datasets/1.1.0/atac_pbmc_10k_nextgem; access date: 01/2021).

#### Single-cell ATAC-seq data processing

For the single-cell ATAC-seq data processing, we employed both Cell Ranger ATAC 1.1.0 (https://support.10xgenomics.com/single-cell-atac/software/downloads/latest) and snapATAC pipelines^57^. Specifically, we first used *count* operation of the Cell Ranger ATAC pipeline to perform a quality assessment, preprocessing, and read alignment, yielding position-sorted barcoded and read-filtered BAM files. We then selected cells with 0.15 < FRiP < 0.5 and the number of reads with uniquely mapped identifiers (UMI) >10,000. Cells across from all datasets were harmonized by Harmony [ref] and clustered based on a K-nearest neighbor (KNN) algorithm with a Louvain community detection (# of eigen dimensions = 47). Consequently, we derived unsupervised snATAC-seq cell clusters enriched to known 16 kidney cell types (**Fig. S7a**) and 12 PBMC cell types (**Fig. S8a**). Specifically, we annotated cell-type of the unsupervised snATAC-seq clusters by label transfer from the cluster of snRNA-seq data that have differential gene expression of known cell-markers. Label transfer is based on correlating open-chromatin activities of gene bodies and mRNA expressions transcribed from the corresponding genes. To visualize the cell clusters, we used Uniform Manifold Approximation and Projection for Dimension Reduction (UMAP)^58^.

#### Subsampling of cells and cross-validation of the optimized peaks

To simulate rare cell types, we subsampled cells with n=200, 500, and 1000 from each of the five abundant cell types in the kidney and PBMC snATAC-seq datasets: PT (n≈8,000), LH (n≈3,800), DCT (n≈1,800), CD4M (n≈1,800), CD14M (n≈4,500). We called peaks using reads aggregated across the same cell type (i.e., pseudo-bulk) from the subsampled cells and conducted the gkmQC peak-calling optimization. We then compared the peaks recovered by the optimization to the peaks called from the entire cells. To quantify the degree to which the optimization process recovers true peaks, we calculated the fraction of the added peaks overlapping original peaks from entire cells.

## Code availability

gkmQC is available in https://github.com/Dongwon-Lee/gkmQC.

IPython notebooks to reproduce the results are available in https://github.com/Dongwon-Lee/gkmQC-manuscript.

## Acknowledgements

We thank all the members of the Sampson laboratory for their helpful discussion. The research reported here was supported by NIH grants HL086694 and HL141980, Boston Children’s Hospital OFD/BTREC/CTREC Faculty Development Fellowship Award, and a Manton Center Endowed Scholar Award. MGS is supported by NIH RO1DK119380 and RC2DK122397. We would like to acknowledge Boston Children’s Hospital’s High-Performance Computing Resources BCH HPC Clusters Enkefalos 2 (E2) made available for conducting the research. Software used in the project was installed and configured by BioGrids^59^.

## Author contribution

S.K.H. and D.L. conceived of the study; S.K.H performed all the analyses; S.K.H., D.L., A.C., and M.G.S. contributed to methods development; Y.M., P.C.W., and B.D.H., contributed to the analyses of kidney snATAC-seq; all authors were involved in manuscript writing and revisions.

