## Supplemental Data for "Quality assessment and refinement of chromatin accessibility data using a sequence-based predictive model"

### Supplementary Figures and legends

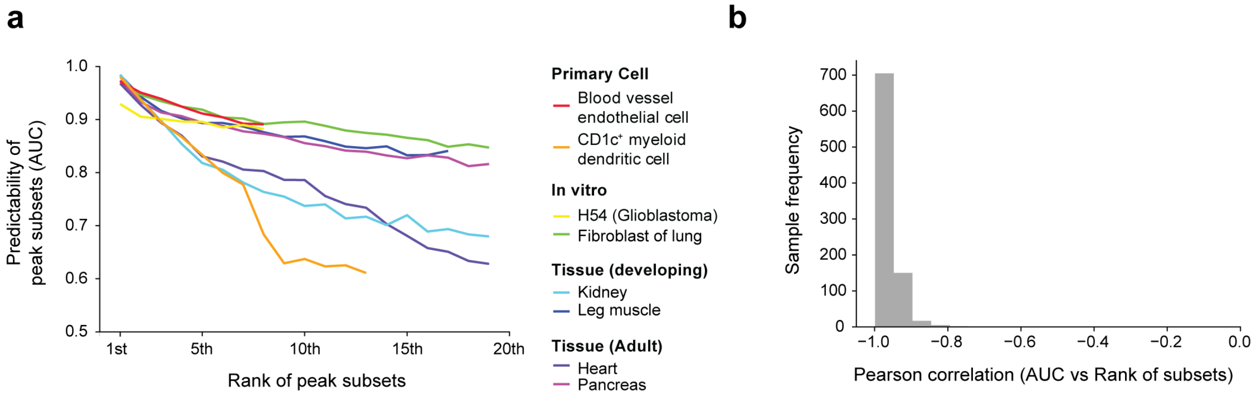


**Figure S1. Correlation between predictability and signal strength of peak subsets. (a)** A plot of predictability (AUC, Y-axis) and the rank of signal strengths of peak subsets (X-axis). Each line represents a distinct sample. A total of 8 representative examples including primary/in vitro differentiated cells and developing/adult tissues are presented. **(b)** The distribution of Pearson correlations between peak predictability and signal strength across ENCODE datasets; correlations between AUC and the rank of signal strengths were computed per sample.

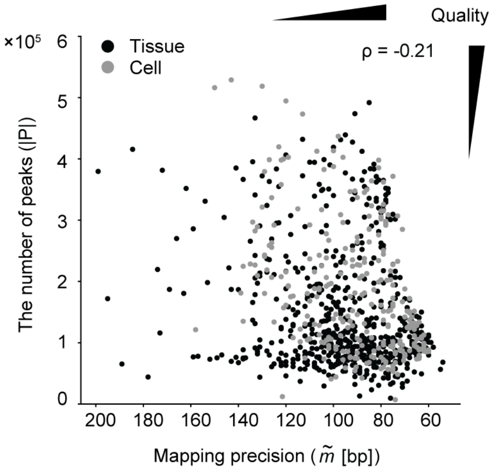


**Figure S2. Comparison of two quality parameters ($\tilde{m}$ and $|P|$).** A weak anti-correlation between the precision of peak location and the number of peaks demonstrates a trade-off between two different quality parameters.

a

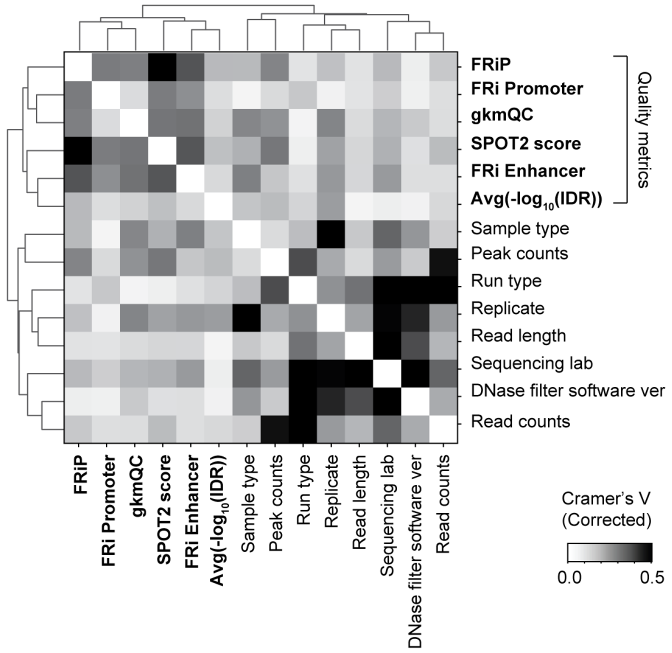
FRiP
FRi Promoter
gkmQC
SPOT2 score
FRi Enhancer
Avg(-log10(IDR))
Sample type
Peak counts
Run type
Replicate
Read length
Sequencing lab
DNase filter software ver
Read counts
Quality metrics
Cramer's V (Corrected)
0.0
0.5

b

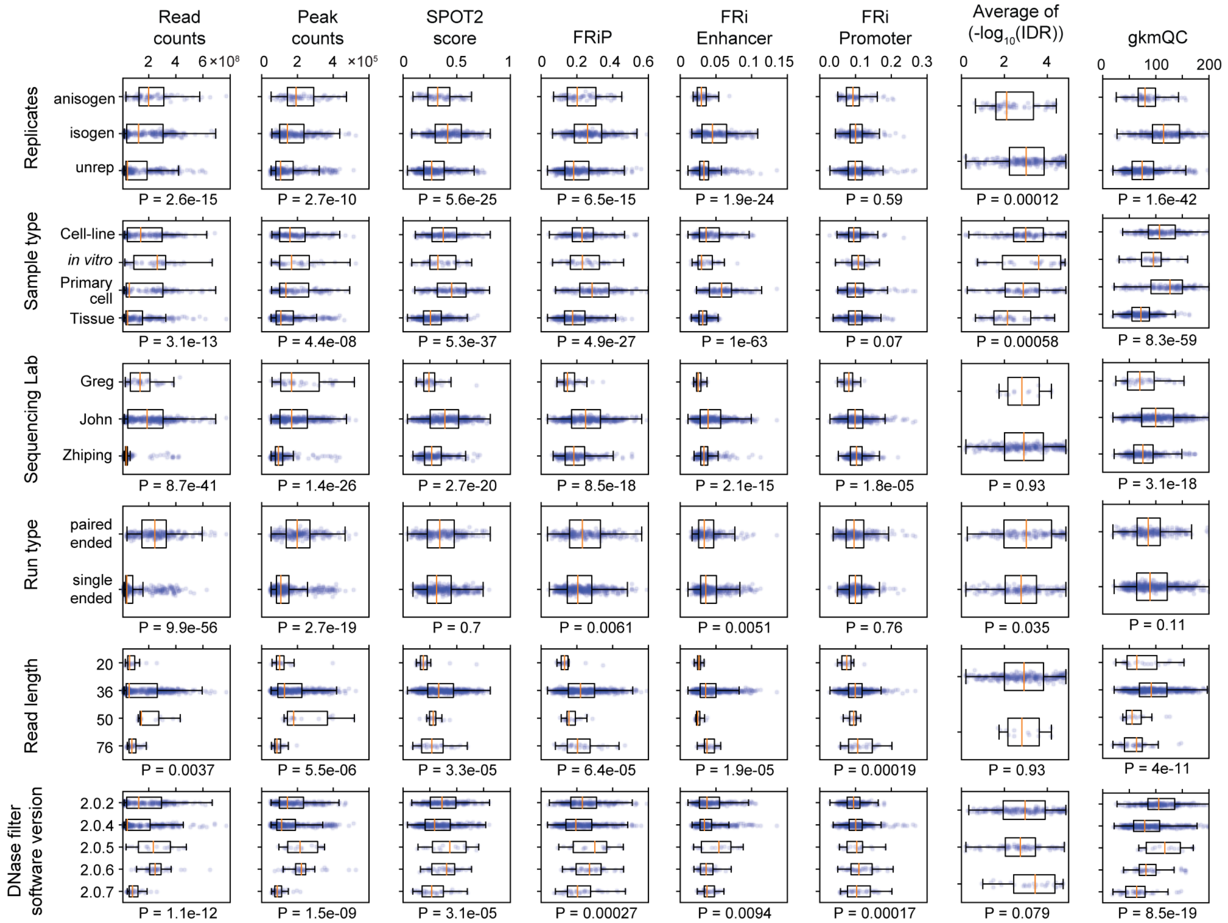
Read counts
Peak counts
SPOT2 score
FRiP
FRi Enhancer
FRi Promoter
Average of (-log10(IDR))
gkmQC
Replicates
anisogen
isogen
unrep
P = 2.6e-15
P = 2.7e-10
P = 5.6e-25
P = 6.5e-15
P = 1.9e-24
P = 0.59
P = 0.00012
P = 1.6e-42
Sample type
Cell-line
in vitro
Primary cell
Tissue
P = 3.1e-13
P = 4.4e-08
P = 5.3e-37
P = 4.9e-27
P = 1e-63
P = 0.07
P = 0.00058
P = 8.3e-59
Sequencing Lab
Greg
John
Zhiping
P = 8.7e-41
P = 1.4e-26
P = 2.7e-20
P = 8.5e-18
P = 2.1e-15
P = 1.8e-05
P = 0.93
P = 3.1e-18
Run type
paired ended
single ended
P = 9.9e-56
P = 2.7e-19
P = 0.7
P = 0.0061
P = 0.0051
P = 0.76
P = 0.035
P = 0.11
Read length
20
36
50
76
P = 0.0037
P = 5.5e-06
P = 3.3e-05
P = 6.4e-05
P = 1.9e-05
P = 0.00019
P = 0.93
P = 4e-11
DNase filter software version
2.0.2
2.0.4
2.0.5
2.0.6
2.0.7
P = 1.1e-12
P = 1.5e-09
P = 3.1e-05
P = 0.00027
P = 0.0094
P = 0.00017
P = 0.079
P = 8.5e-19

**Figure S3**. **Analysis of biological and technical factors affecting quality metrics (a)** Heatmap shows covariation between several technical factors and quality metrics. Cramer's V was used to quantify correlations of continuous and discrete variables. Technical factors were not clustered with quality metrics (bold black). **(b)** Boxplots show differences in quality metrics (x-axis) for several different technical factors. *P*-values were calculated with one-way ANOVA of the quality metric scores with respect to the technical factors.

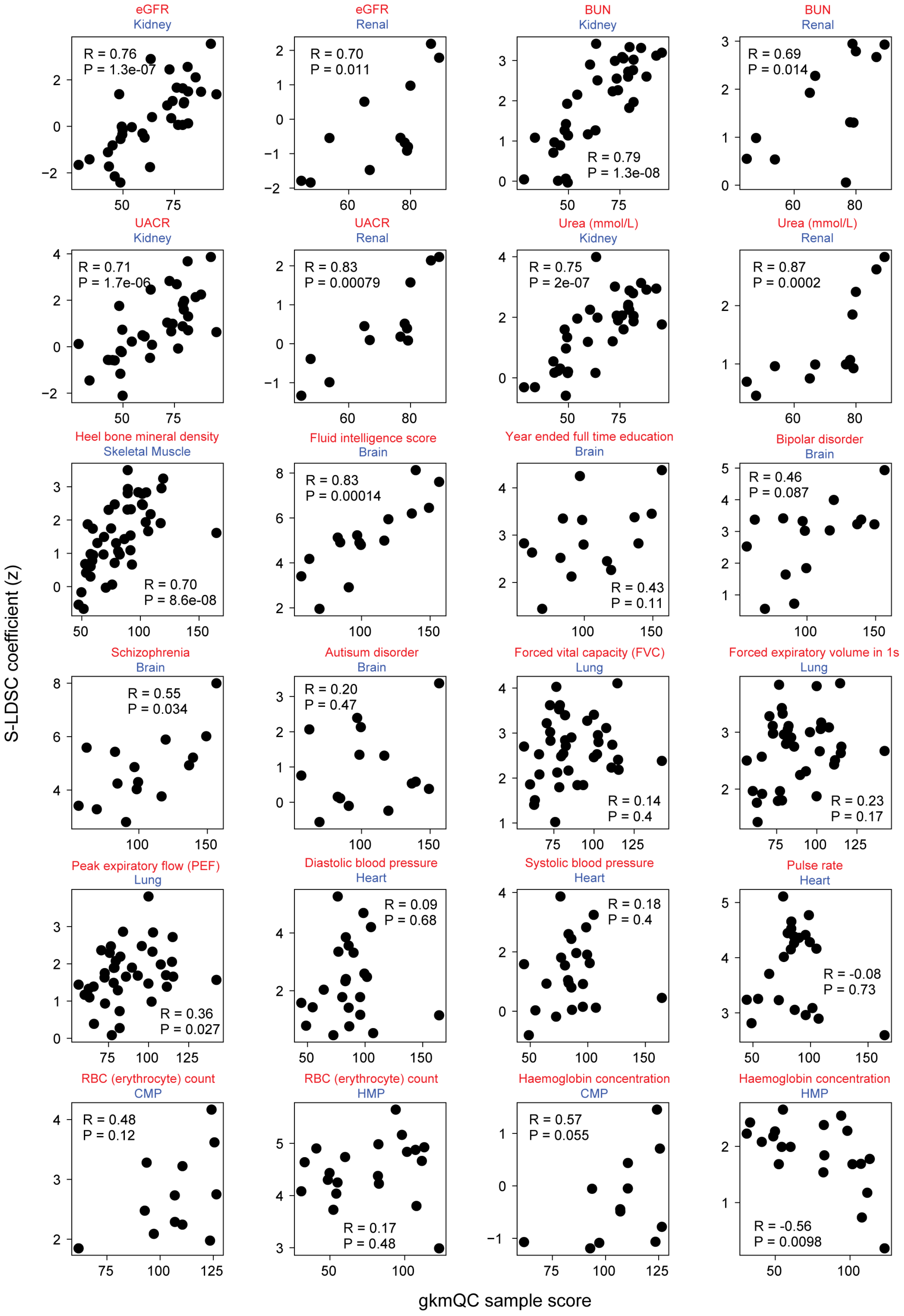

eGFR
Kidney
R = 0.76
P = 1.3e-07
eGFR
Renal
R = 0.70
P = 0.011
BUN
Kidney
R = 0.79
P = 1.3e-08
BUN
Renal
R = 0.69
P = 0.014
UACR
Kidney
R = 0.71
P = 1.7e-06
UACR
Renal
R = 0.83
P = 0.00079
Urea (mmol/L)
Kidney
R = 0.75
P = 2e-07
Urea (mmol/L)
Renal
R = 0.87
P = 0.0002
Heel bone mineral density
Skeletal Muscle
R = 0.70
P = 8.6e-08
Fluid intelligence score
Brain
R = 0.83
P = 0.00014
Year ended full time education
Brain
R = 0.43
P = 0.11
Bipolar disorder
Brain
R = 0.46
P = 0.087
Schizophrenia
Brain
R = 0.55
P = 0.034
Autisum disorder
Brain
R = 0.20
P = 0.47
Forced vital capacity (FVC)
Lung
R = 0.14
P = 0.4
Forced expiratory volume in 1s
Lung
R = 0.23
P = 0.17
Peak expiratory flow (PEF)
Lung
R = 0.36
P = 0.027
Diastolic blood pressure
Heart
R = 0.09
P = 0.68
Systolic blood pressure
Heart
R = 0.18
P = 0.4
Pulse rate
Heart
R = -0.08
P = 0.73
RBC (erythrocyte) count
CMP
R = 0.48
P = 0.12
RBC (erythrocyte) count
HMP
R = 0.17
P = 0.48
Haemoglobin concentration
CMP
R = 0.57
P = 0.055
Haemoglobin concentration
HMP
R = -0.56
P = 0.0098
S-LDSC coefficient (z)
gkmQC sample score

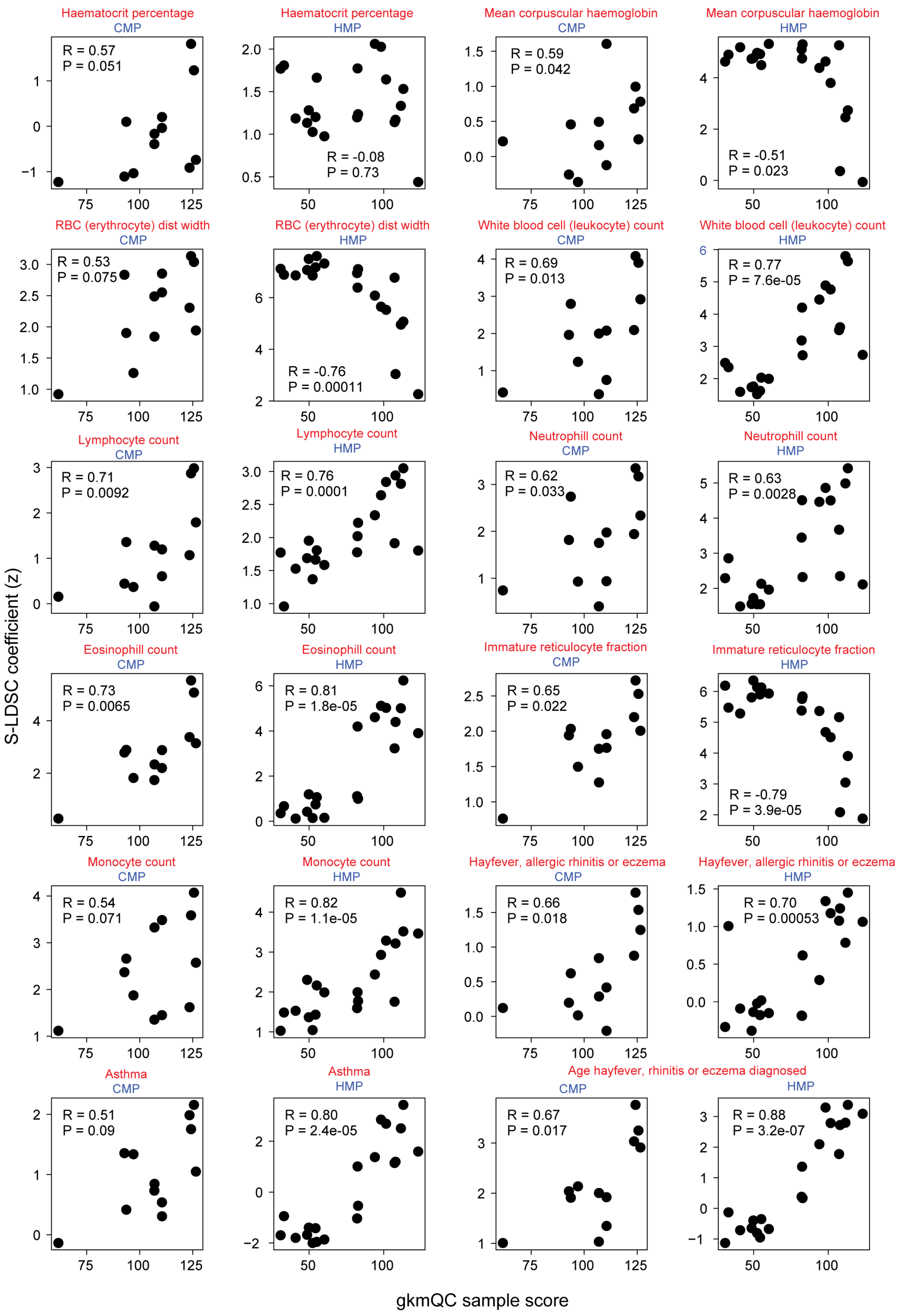
S-LDSC coefficient (z)
gkmQC sample score
Haematocrit percentage
CMP
R = 0.57
P = 0.051
Haematocrit percentage
HMP
R = -0.08
P = 0.73
Mean corpuscular haemoglobin
CMP
R = 0.59
P = 0.042
Mean corpuscular haemoglobin
HMP
R = -0.51
P = 0.023
RBC (erythrocyte) dist width
CMP
R = 0.53
P = 0.075
RBC (erythrocyte) dist width
HMP
R = -0.76
P = 0.00011
White blood cell (leukocyte) count
CMP
R = 0.69
P = 0.013
White blood cell (leukocyte) count
HMP
R = 0.77
P = 7.6e-05
Lymphocyte count
CMP
R = 0.71
P = 0.0092
Lymphocyte count
HMP
R = 0.76
P = 0.0001
Neutrophill count
CMP
R = 0.62
P = 0.033
Neutrophill count
HMP
R = 0.63
P = 0.0028
Eosinophill count
CMP
R = 0.73
P = 0.0065
Eosinophill count
HMP
R = 0.81
P = 1.8e-05
Immature reticulocyte fraction
CMP
R = 0.65
P = 0.022
Immature reticulocyte fraction
HMP
R = -0.79
P = 3.9e-05
Monocyte count
CMP
R = 0.54
P = 0.071
Monocyte count
HMP
R = 0.82
P = 1.1e-05
Hayfever, allergic rhinitis or eczema
CMP
R = 0.66
P = 0.018
Hayfever, allergic rhinitis or eczema
HMP
R = 0.70
P = 0.00053
Asthma
CMP
R = 0.51
P = 0.09
Asthma
HMP
R = 0.80
P = 2.4e-05
Age hayfever, rhinitis or eczema diagnosed
CMP
R = 0.67
P = 0.017
HMP
R = 0.88
P = 3.2e-07

**Figure S4. Correlation analyses of gkmQC sample scores and S-LDSC coefficients for 48 pairs of relevant tissues and phenotypes (2 pages).** Dots in the scatterplots represent samples of chromatin accessibility data. The X-axis is the gkmQC sample scores, and the Y-axis is the S-LDSC coefficient. Correlations are Pearson's correlation coefficients. The title of each scatterplot shows a tissue- or cell-type for chromatin-accessibility data (Blue) and a relevant GWAS trait (Red).

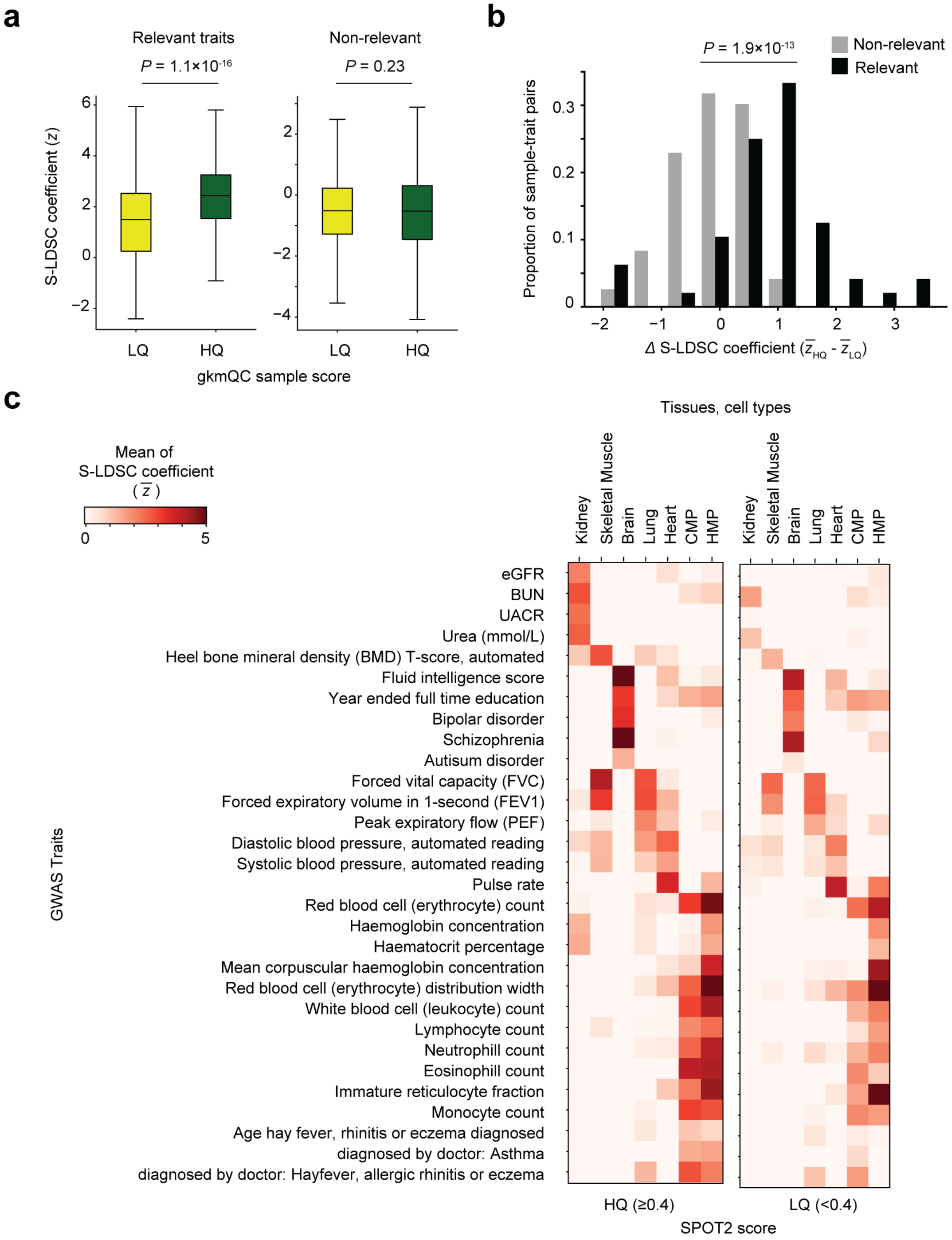


**Figure S5**. **Contribution of tissue-specific peaks in high-quality samples to relevant traits. (a)** Boxplots show distributions of S-LDSC coefficients of high- and low-quality samples paired with relevant (left) and non-relevant traits (right). Mann-Whitney *U* test is used to test the significance of the differences. **(b)** Histograms show differences in mean S-LDSC coefficients between high- and low-quality samples for relevant and non-relevant traits. Paired *t*-test is used to test the significance of the differences. **(c)** the same analysis as **Figure 4b** was repeated using SPOT2 as a quality metric. High-quality samples are SPOT2 scores >0.4 based on ENCODE experimental guideline (https://www.encodeproject.org/data-standards/dnase-seq/). Renal cortex tissue is excluded due to lack of high-quality samples with SPOT2 >0.4.

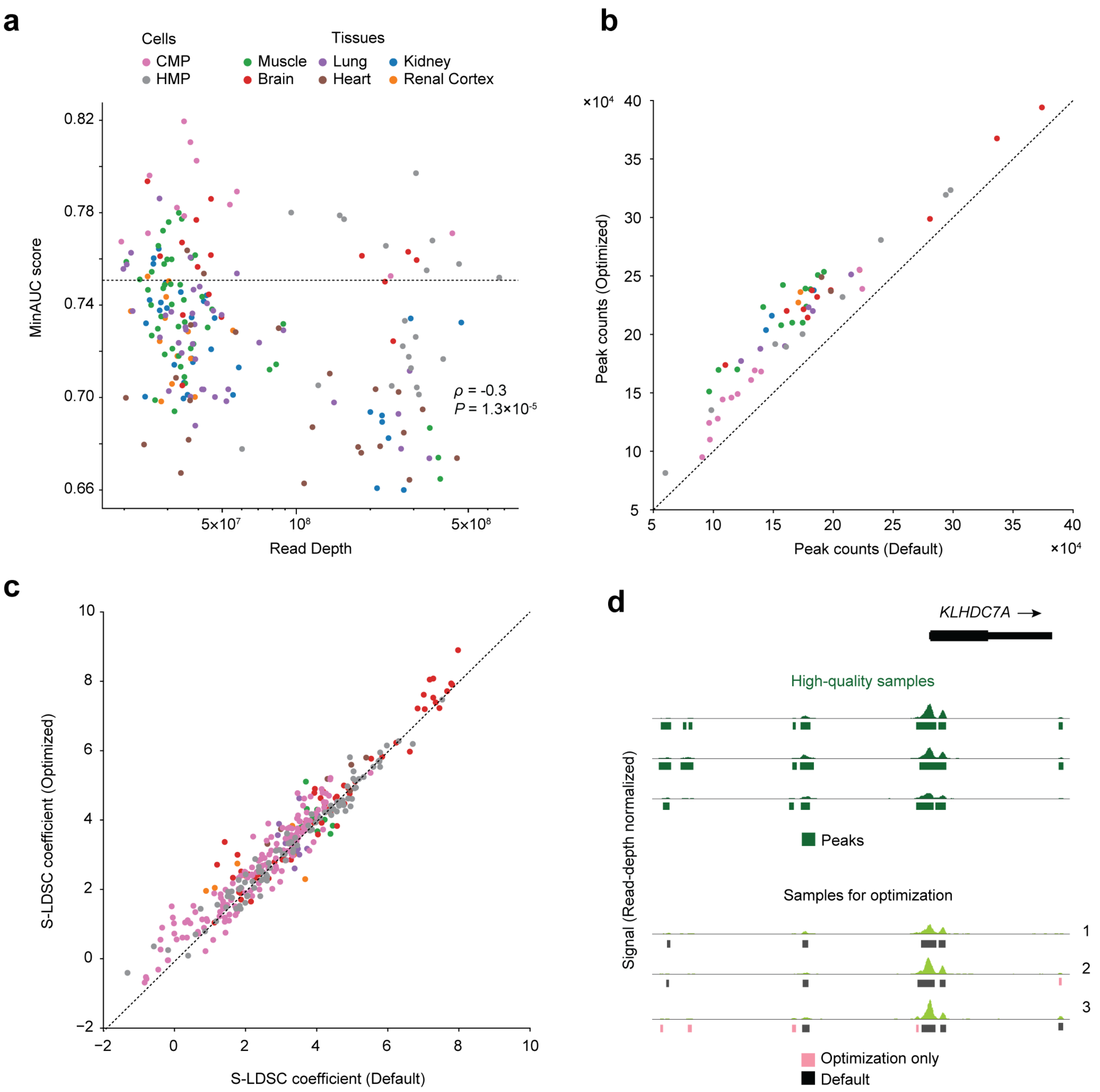


**Figure S6**. **Optimizing peak-calling from bulk chromatin accessibility data. (a)** Scatterplot comparing read depth (total read counts) with MinAUC using Spearman rank correlation coefficients. Each dot is a distinct sample with colors representing tissues and cell types. **(b)** Peak counts before and after gkmQC optimization. **(c)** S-LDSC coefficients from the heritability analysis of relevant GWAS traits before and after optimization. **(d)**. Newly found peaks after optimization recapitulate peaks observed in high-quality samples at the *KLHDC7A* locus (**Fig. 2b**).

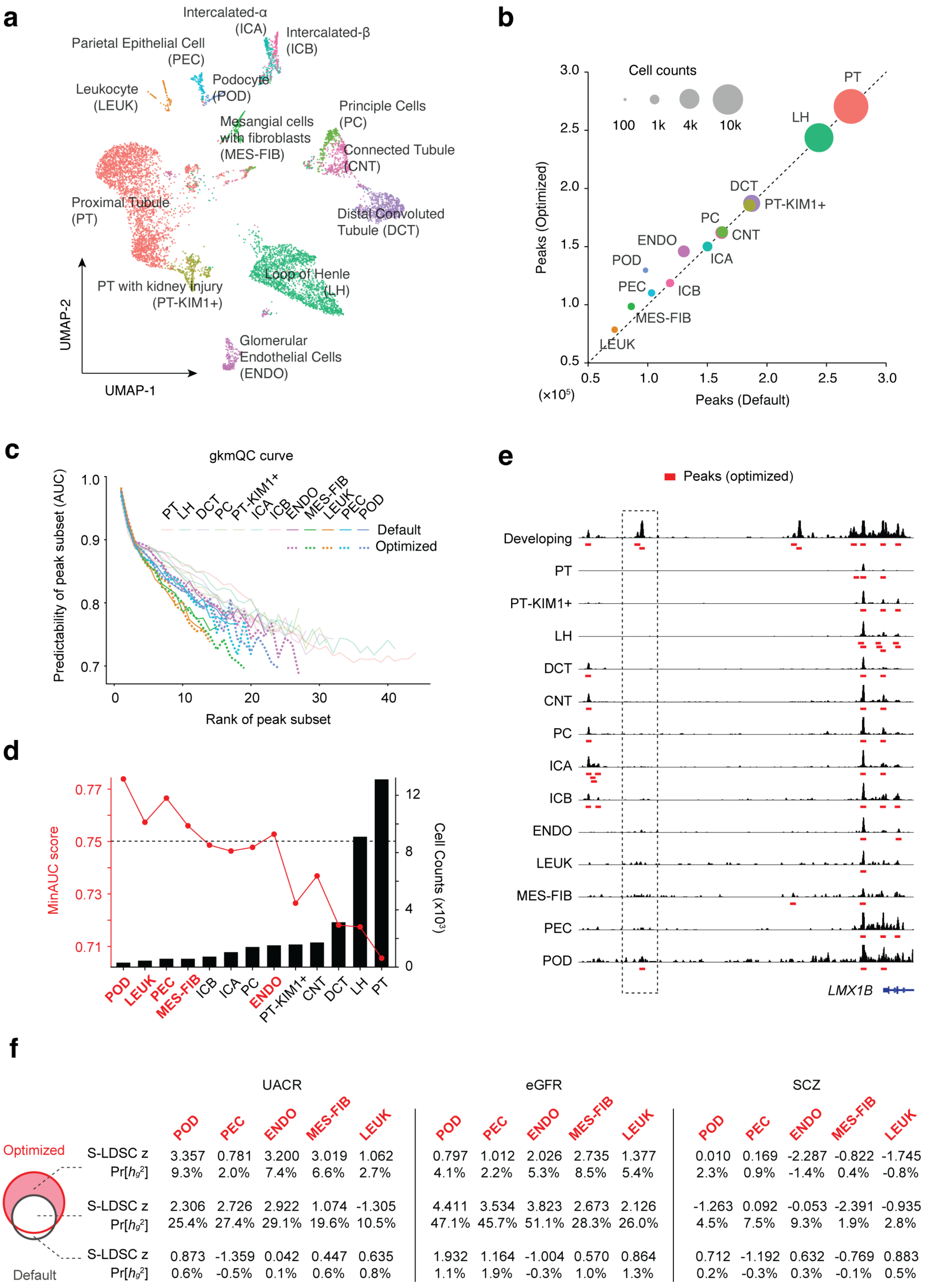
a
Intercalated-α (ICA)
Intercalated-β (ICB)
Parietal Epithelial Cell (PEC)
Podocyte (POD)
Leukocyte (LEUK)
Mesangial cells with fibroblasts (MES-FIB)
Principle Cells (PC)
Connected Tubule (CNT)
Proximal Tubule (PT)
Distal Convoluted Tubule (DCT)
Loop of Henle (LH)
PT with kidney injury (PT-KIM1+)
Glomerular Endothelial Cells (ENDO)
UMAP-2
UMAP-1
b
Cell counts
100
1k
4k
10k
Peaks (Optimized)
Peaks (Default)
(×10⁵)
PT
LH
DCT
PT-KIM1+
PC
CNT
ENDO
ICA
POD
PEC
ICB
MES-FIB
LEUK
c
gkmQC curve
Predictability of peak subset (AUC)
Rank of peak subset
PT LH DCT PC PT-KIM1+ ICA ICB ENDO MES-FIB LEUK PEC POD
Default
Optimized
d
MinAUC score
Cell Counts (x10³)
POD LEUK PEC MES-FIB ICB ICA PC ENDO PT-KIM1+ CNT DCT LH PT
e
Peaks (optimized)
Developing
PT
PT-KIM1+
LH
DCT
CNT
PC
ICA
ICB
ENDO
LEUK
MES-FIB
PEC
POD
LMX1B
f
UACR
eGFR
SCZ
POD PEC ENDO MES-FIB LEUK
Optimized
Default
S-LDSC z 3.357 0.781 3.200 3.019 1.062 | 0.797 1.012 2.026 2.735 1.377 | 0.010 0.169 -2.287 -0.822 -1.745
Pr[h_g²] 9.3% 2.0% 7.4% 6.6% 2.7% | 4.1% 2.2% 5.3% 8.5% 5.4% | 2.3% 0.9% -1.4% 0.4% -0.8%
S-LDSC z 2.306 2.726 2.922 1.074 -1.305 | 4.411 3.534 3.823 2.673 2.126 | -1.263 0.092 -0.053 -2.391 -0.935
Pr[h_g²] 25.4% 27.4% 29.1% 19.6% 10.5% | 47.1% 45.7% 51.1% 28.3% 26.0% | 4.5% 7.5% 9.3% 1.9% 2.8%
S-LDSC z 0.873 -1.359 0.042 0.447 0.635 | 1.932 1.164 -1.004 0.570 0.864 | 0.712 -1.192 0.632 -0.769 0.883
Pr[h_g²] 0.6% -0.5% 0.1% 0.6% 0.8% | 1.1% 1.9% -0.3% 1.0% 1.3% | 0.2% -0.3% 0.3% -0.1% 0.5%

**Figure S7**. **Peak-calling optimization of kidney snATAC-seq data identifies more functional peaks for rare cell types. (a)** UMAP plot of kidney snATAC-seq data. Color is based on annotation of known kidney cell types. **(b)** Comparison of peak counts before and after optimization where each dot represents kidney cell type in (a). Dot sizes represent cell counts of the corresponding cell types. **(c)** gkmQC curves for peaks from pseudo-bulk reads of kidney cells. Dashed lines are gkmQC curves for optimized peak-calling. The five cell types with MinAUC >0.75 were optimized. **(d)** MinAUC scores of kidney cell types (red line with dots; left Y-axis) are anti-correlated with cell counts (black bars; right Y-axis), demonstrating more significant optimization in rarer cell types; cell types with MinAUC >0.75 are highlighted (red). **(e)** A representative locus upstream of *LMX1B* containing a podocyte-specific peak. All kidney cell types in the snATAC-seq and developing (DNase-seq) are shown. **(f)** Heritability is compared between optimized and default peaks for five rare cell types with MinAUC >0.75. Similar to **Fig. 5d**, the table presents heritability for three disjoint peak subsets; optimization-only (top), commonly found before/after optimization (middle), and only with default values (bottom).

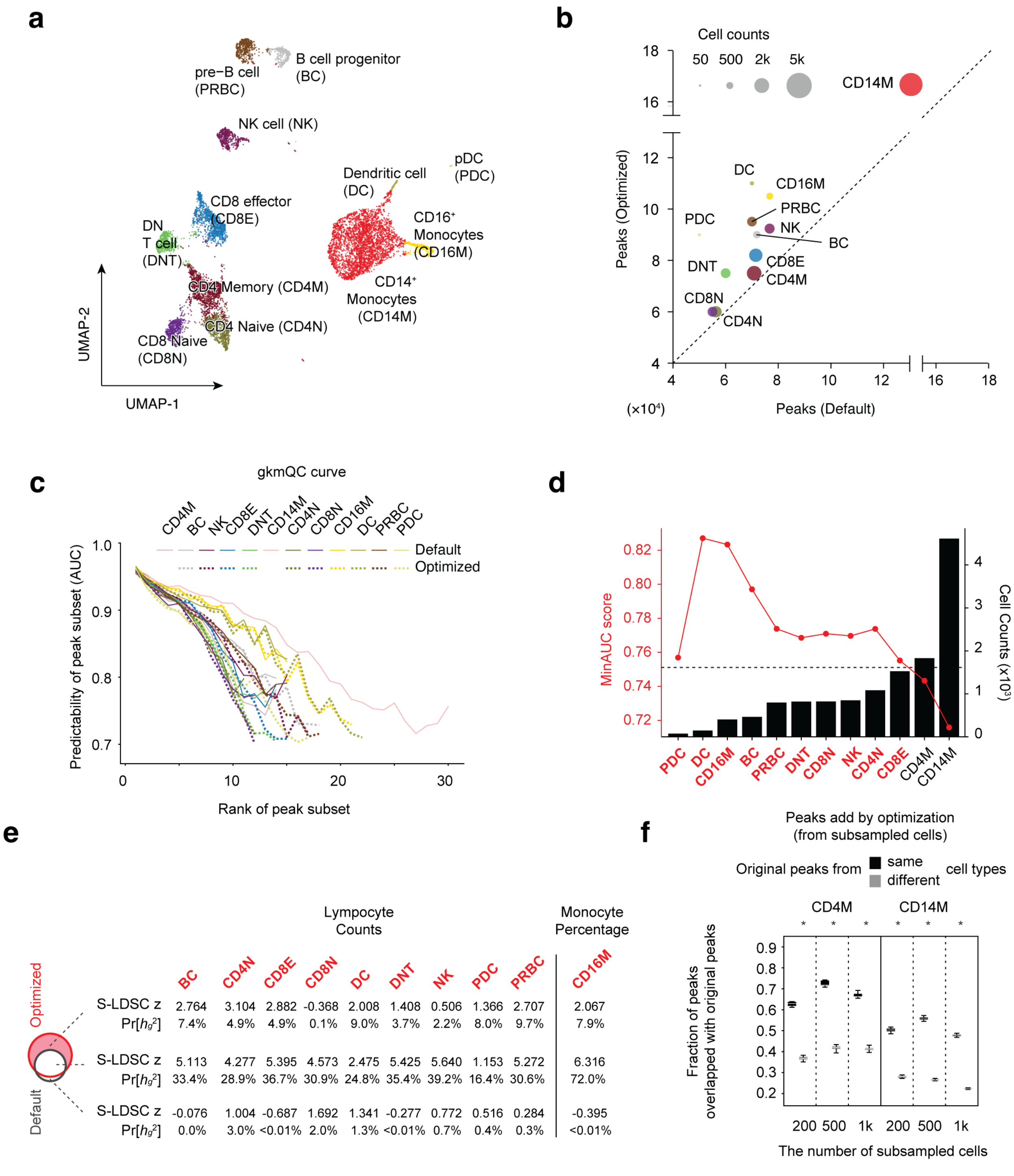


**Figure S8**. **Peak-calling optimization of PBMC snATAC-seq data.** Figures **a-e** are analogous to **Figures S7a-d** and **f**, except that ten cell types were optimized for PBMC snATAC-seq data. Figure **f** is analogous to **Figure 6d**. Here, two major cell types (CD14+ Monocyte and CD4+ Memory T cells) were analyzed.

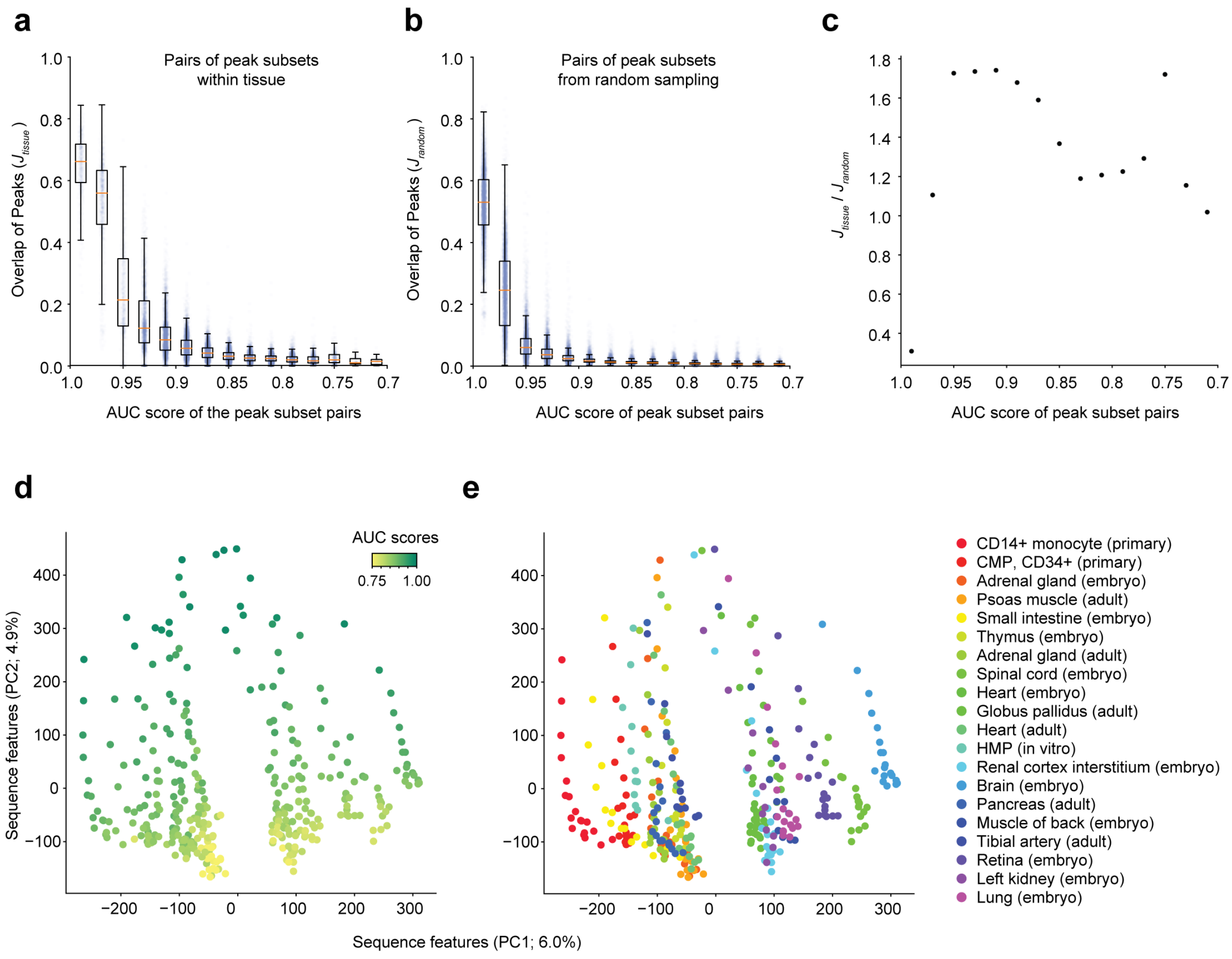


**Figure S9**. **Differences in peak-calling signals between peak subsets reflect differences in tissue-specificity.** Figures a-c depict relative degrees of tissue-specificity of peak subsets as a function of their peak predictability (AUC). We calculated overlaps of peak-subset pairs in a similar AUC range **(a)** from the same tissue and **(b)** from different tissues via random sampling. Jaccard index coefficients were used to quantify overlap. **(c)** The overlap ratios between (a) and (b) ($J_{tissue}$ / $J_{random}$) are calculated for each of the AUC ranges. (**d and e**) Principal component 1 (PC1) and PC2 from PCA analysis of the trained sequence features are shown for several different tissues. Peak subsets are represented as dots, color-coded for **(d)** AUCs and **(e)** tissues. Peak subsets in a medium range of predictability scores (0.8< AUC <0.95) have more tissue-specific sequence features.

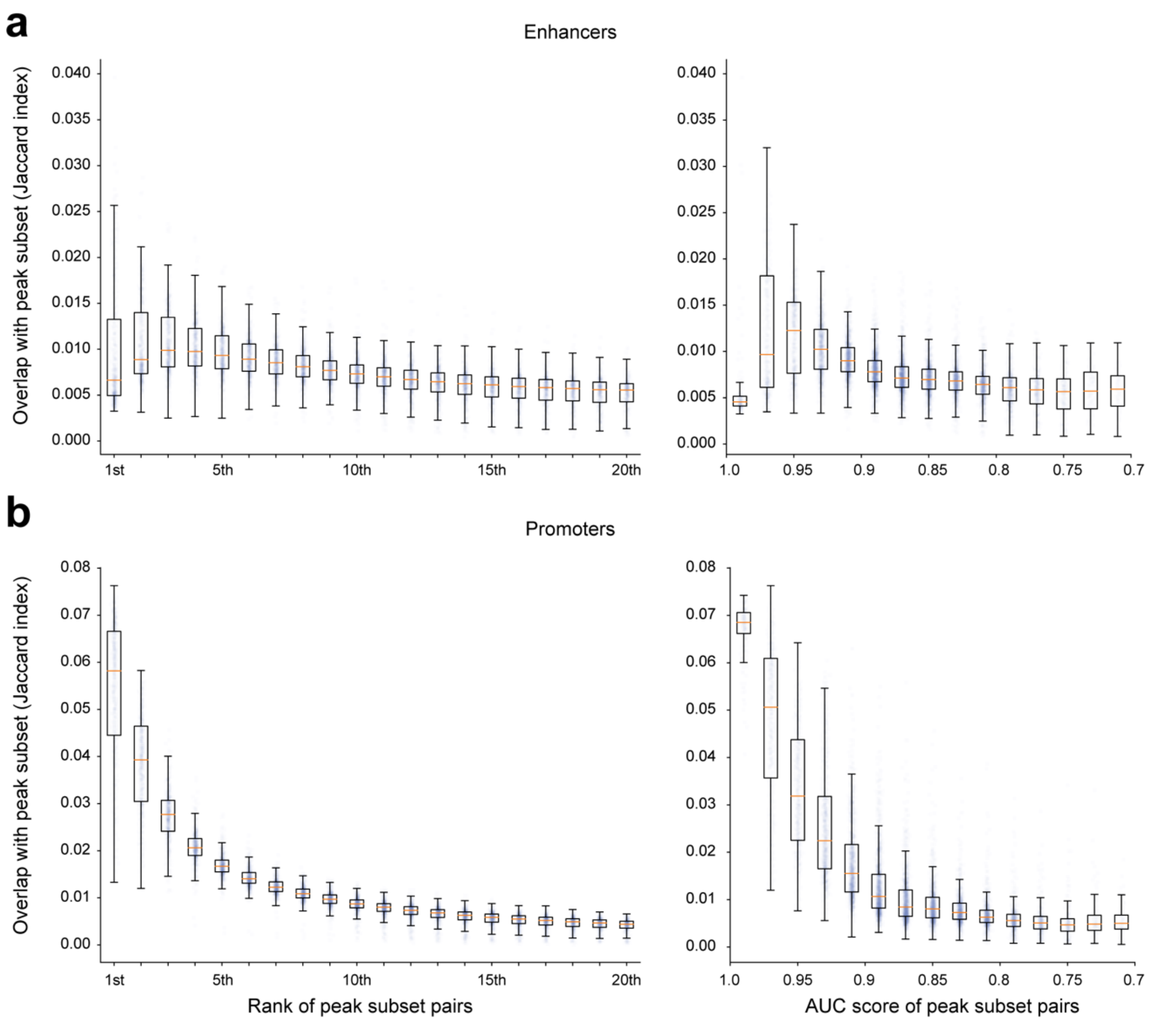


**Figure S10**. **Enrichment analysis of enhancers and promoters with respect to peak signal and predictability. (a)** FANTOM5 (distal) enhancers and **(b)** promoters are compared to peak subsets according to their ranks and AUCs. Overlaps between two peak sets are calculated by the Jaccard index. Boxplots represent overlap distributions across different samples.

### Supplementary Tables and legends

**Table S1.** Metadata and quality metric statistics of 886 ENCODE DNase-seq samples.

**Table S2**. Results from partitioned heritability analysis using 200 ENCODE DNase-seq samples and relevant GWAS traits.

**Data S1.** Archived files including the BED files of optimized peaks for 58 DNase-seq data.

**Data S2.** Archived files including the BED files of optimized peaks for kidney and PBMC snATAC-seq data
